# Cortical organoid diffusion MRI reveals protocol-associated spatial reproducibility and increasing apparent diffusivity with diffusion time

**DOI:** 10.64898/2026.09.29.755111

**Authors:** Andrés le Boeuf Fló, Ekin Taskin, Oriana Lavielle, Athanasios Grigoriou, Katarzyna Pierzchała, Thanh Phong Lê, Ileana Jelescu, Zeljka Krsnik, Théo Ribierre, Jean-Philippe Thiran, Jonathan Rafael-Patiño, Erick Jorge Canales-Rodríguez, Elda Fischi-Gómez

## Abstract

Early human cortical development involves rapid cytoarchitectural changes that are difficult to study non-invasively. Diffusion MRI can probe developing microstructure and membrane water exchange, but model assumptions are difficult to validate in human tissue. Cortical organoids provide a tractable platform for relating diffusion measurements to tissue architecture. We scanned six fixed cortical organoids at 9.4 T, three grown under directed and three under undirected differentiation protocols, at three diffusion times and b-values up to 7000 s/mm^2^. Center-to-periphery profiles of diffusion kurtosis imaging (DKI) metrics were reproducible across directed organoids but more variable across undirected organoids, consistent with nuclear staining. Unexpectedly, DKI-derived apparent mean diffusivity (MD) increased with diffusion time in more than 80% of tissue voxels in both batches, opposite to previous organoid measurements. Across 12,348 analytically simulated two-compartment substrates and Monte Carlo diffusion simulations in sphere packings, this trend emerged in permeable substrates with a marked contrast between extracellular and apparent intracellular diffusivities. Detailed Monte Carlo trajectory analysis of a representative substrate showed that fitted MD increased despite decreases in displacement-derived and second-cumulant diffusivities, revealing an exchange-sensitive finite-b effect involving higher-order signal contributions. These findings support using cortical organoids to probe early microstructure, test diffusion models, and investigate potential emerging diffusion biomarkers.

## Introduction

Early human cortical development involves rapid changes in cellular composition, laminar organization, and tissue architecture that are difficult to characterize non-invasively ^1;2^. Diffusion magnetic resonance imaging, and particularly biophysical models of gray-matter diffusion, offer a potential route to infer microstructural features such as soma and neurite fractions, compartmental diffusivities, and water exchange across cell membranes ^3–5^. However, these estimates rely on assumptions about tissue geometry and permeability that remain difficult to validate directly ^6^. Histological validation relies largely on animal or postmortem tissue and is especially constrained for scarce developing human cortex, where postmortem delay, fixation, and limited tissue availability further complicate interpretation ^7–9^. Physical diffusion phantoms can reproduce selected geometric or exchange properties, but do not capture the coupled cellular organization and permeability of developing gray matter ^10;11^. More generally, microstructural parameters used to interpret gray matter in vivo are inappropriate for the developing cortex.

Human cortical organoids offer an alternative to physical phantoms. They self-organize into layered, cell-dense tissue that recapitulates early corticogenesis ^12–14^, and can be imaged and then stained on the same specimen. Undirected protocols rely on intrinsic self-organization ^14^. In contrast, directed protocols guide differentiation toward a defined region such as the dorsal forebrain ^15^ or other brain regions of interest ^16–20^, and produce a reproducible diversity of cortical cell types and developmental trajectories, generating cortical tissue more reproducibly across organoids ^21^. However, their characterization relies mainly on optical microscopy, which is limited by penetration depth unless samples are cleared, and on electron microscopy, which is destructive and restricted to small volumes ^22^.

Magnetic resonance imaging (MRI) avoids these constraints: it images the whole organoid non-invasively. Several studies have shown that MR contrast tracks cytoarchitecture: T_1_-weighted imaging at 3T separates regions of high and low cellularity in bioink-embedded cortical organoids, validated against nuclear staining of the same samples ^23^; 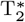 -weighted imaging at 9.4 T resolves cystic from compact tissue and supports longitudinal monitoring over two months ^24^. Diffusion MRI (dMRI) adds sensitivity below the voxel resolution level, and a few studies have applied it to organoids. Multishell imaging of fixed hippocampal organoids at 11.7 T resolved rosette-like structures as isotropic cores surrounded by anisotropic tissue matching immunostained neuronal processes in different organoids ^25^; Diffusion Tensor Imaging (DTI) at 9.4 T found higher diffusivity in cystic than in compact tissue ^24^; Apparent Diffusion Coefficient (ADC) mapping at 14 T separated fast- and slow-diffusing regions within individual fixed organoids ^26^; and ADC at 15.2 T detected elevated water diffusion in a *LIS1* lissencephaly model that reverted after enzymatic digestion of the extracellular matrix ^27^. More recently, ultra-high-field dMRI at 28.2 T extended living-organoid measurements to higher spatial resolution and richer diffusion encoding, reporting anisotropy, maturation-dependent differences in ADC and kurtosis, and correspondence with three-dimensional light-sheet microscopy ^28^.

These studies leave two complementary gaps. First, diffusion-time dependence remains poorly characterized in organoids. Most studies have used a single diffusion time, making diffusivity sensitive to a combination of compartment size, intrinsic diffusivity, and exchange. To our knowledge, only two studies have systematically varied diffusion time: in fixed forebrain organoids, diffusivity decreased over t = 7–200 ms, with most of the time dependence occurring below approximately 20 ms ^29^; in living cortical organoids, ADC generally decreased between 2.7 and 25 ms, with a transient increase at the periphery of a young organoid ^28^. Water exchange has also been investigated in cortical organoids through the diffusion-time dependence of kurtosis, using a Kärger-based framework ^29^. Second, although guided and unguided differentiation protocols are known to produce differences in organoid organization and reproducibility, these differences have not been investigated from a dMRI perspective. In particular, it remains unknown whether dMRI can capture protocol-dependent differences in the spatial organization of organoid tissue and whether these spatial diffusion signatures are reproducible across organoids.

Here we scanned six human cortical organoids at 9.4 T: three grown under a directed differentiation protocol and three under an undirected one, fixed and prepared identically, and imaged at three diffusion times across five *b*-shells up to 7000 s/mm^2^ with echo time held constant. We asked two questions. First, does diffusion contrast reflect organoid cytoarchitecture? Using diffusion kurtosis imaging (DKI), we find that center-to-periphery profiles of mean diffusivity, fractional anisotropy, and mean kurtosis are reproducible across the three directed organoids but substantially more variable across the three undirected organoids, revealing batch-associated differences in spatial organization that are consistent with the known differences between the two differentiation strategies. Nuclear staining further shows that the undirected organoids themselves differ in their internal architecture, supporting the biological basis of the observed diffusion variability. We additionally fit the Neurite Exchange Imaging (NEXI) two-compartment model ^3^ to investigate the microstructural and exchange properties underlying these signals. Second, how does diffusion evolve with diffusion time in tissue dominated by cell bodies and short processes? In conventional coarse-grained models, apparent diffusivity is expected to decrease with increasing diffusion time, consistent with previous measurements in organoids ^28;29^. We instead find that the apparent Mean Diffusivity (MD) increases in more than 80% of tissue voxels in both batches, across all six organoids. Across 12,348 two-compartment exchange simulations, no impermeable substrate reproduced this increase within the tested parameter space; the observed behaviour emerged only when membranes were permeable, and the extracellular diffusivity was substantially higher than the apparent intracellular diffusivity. Monte Carlo trajectory analysis further showed that DKI-derived MD increased in a representative simulated substrate despite decreases in displacement-derived and second-cumulant diffusivities, demonstrating that finite-*b* estimates can exhibit a diffusion-time trend opposite to that of the underlying diffusion process.

## Results

Figure 1 summarizes the two organoid batches and the regions used for subsequent analyses. Batch 1 comprised three organoids generated using a directed differentiation protocol, whereas Batch 2 comprised three organoids generated using an undirected protocol.

**Figure 1.**
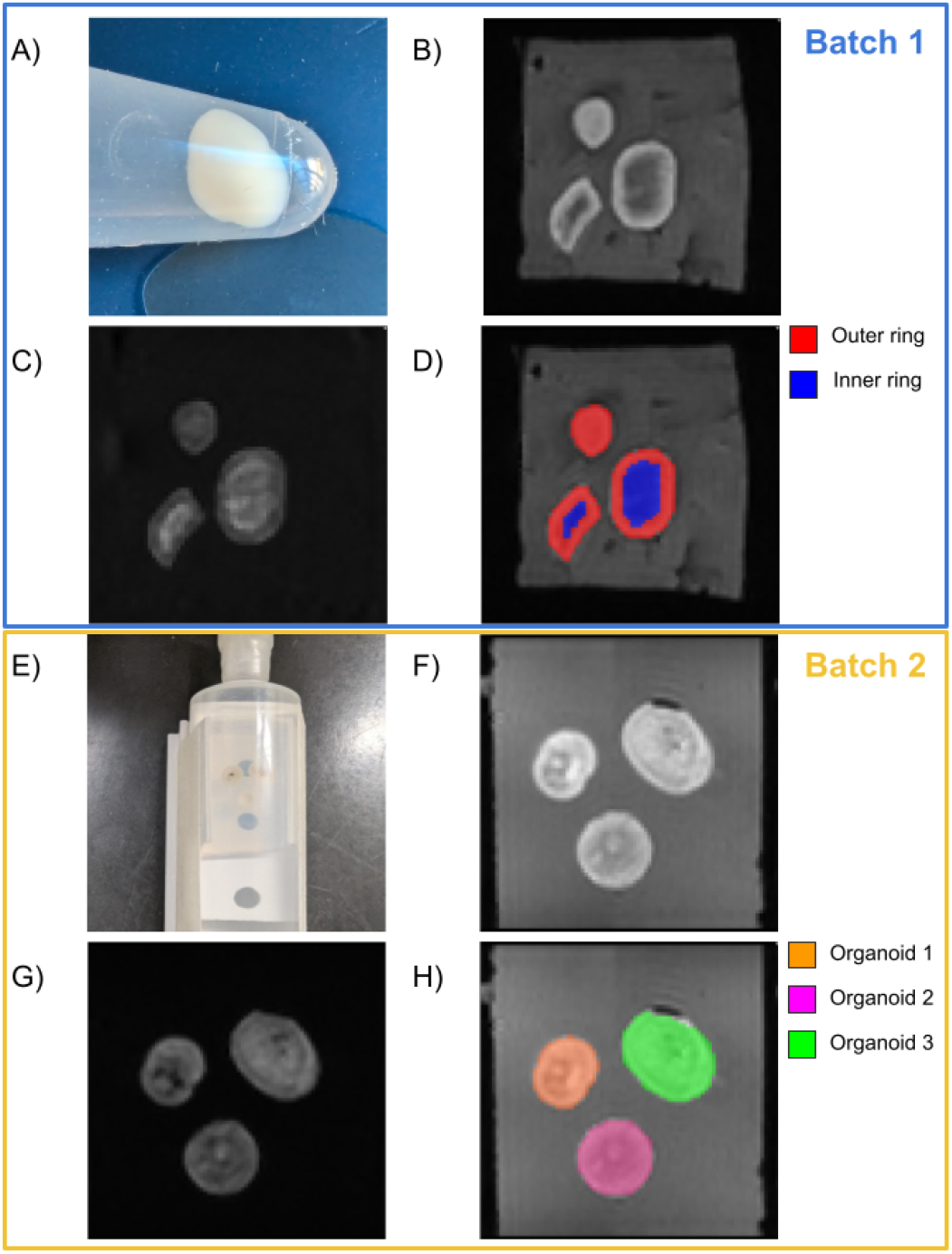
Cortical organoids from Batch 1 (top 4 figures highlighted in blue) and Batch 2 (bottom 4 figures highlighted in yellow). **(A)** Cortical organoid in its test tube; **(E)** Three cortical organoids in their agarose block stabilized within its syringe holder filled with Galden; **(B)** and **(F)** Preprocessed b = 0 images from Δ = 15ms dataset; **(C)** and **(G)** Preprocessed b = 7000 s/mm^2^ images from Δ = 15ms dataset; **(D)** Batch 1 manual segmentation of the inner and outer rings of each organoid overlaid to the b = 0 image; **(H)** Batch 2 manual segmentation of each organoid overlaid to the b = 0 image.

### Directed organoids develop a reproducible radial microstructure that undirected organoids lack

Because the two batches differ by roughly a factor of two in diffusivity, we fitted DKI over batch-specific b-windows (*b* ≤ 3500 s/mm^2^ for Batch 1, *b* ≤ 7000 s/mm^2^ for Batch 2; Methods 4.5, Supplementary Figure S1) and separately at each diffusion time (Δ). All DKI values below follow this convention.

Quantitatively, Batch 1 showed low anisotropy across the sampled diffusion times: Fractional Anisotropy (FA) stays below 0.2 at the 95^th^ percentile and barely moves between Δ = 15 and 38 ms (median 0.093 to 0.097). Batch 2 shows slightly higher FA at short and intermediate diffusion times (median 0.104 and 0.110 at Δ = 15 and 26 ms) that converges towards Batch 1 values by Δ = 38 ms (0.099). Mean Kurtosis (MK) is markedly higher in Batch 2 than in Batch 1 (median 1.71 versus 0.96 at Δ = 15 ms) and decreases with diffusion time in both (Batch 1 median 0.96 → 0.72; Batch 2 1.71 → 1.44).

Two NEXI parameters reached their fitting bounds in a substantial fraction of voxels. The intra-neurite diffusivity (*D*_*i*_) reached its lower bound of 0.1 µm^2^/ms in 42.65% of Batch 1 and 71.87% of Batch 2 voxels, and the exchange time (*t*_*ex*_) reached its upper bound of 150 ms in 0.23% and 22.57% respectively. We report *D*_*i*_ for completeness but do not interpret its precise magnitude or correlate it with the other parameters. Because boundary-hitting indicates limited identifiability and may affect parameter covariance, *D*_*e*_, *f*_*n*_ and *t*_*ex*_ are likewise interpreted as effective, model-dependent estimates rather than independent microscopic measurements. Each parameter statistic is described by batch and region in Table 1.

**Table 1.**
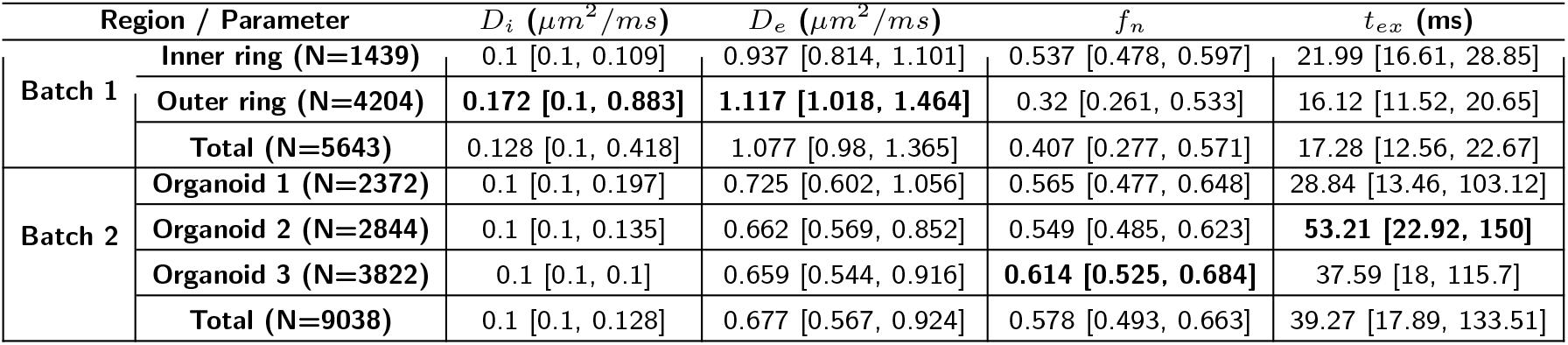
Estimated NEXI parameter values reported as median [first quartile, third quartile]. For Batch 1, regions are subdivided into inner and outer rings, whereas for Batch 2, values are reported separately for each organoid. Bold values indicate the highest estimate of a given parameter within each region across both batches.

In Batch 1, every model parameter separates the inner from the outer ring in the same direction across all three organoids. The NEXI-derived restricted-compartment fraction (*f*_*n*_) is the clearest: *f*_*n*_ reaches a median of 0.54 in the inner ring against 0.32 in the outer ring, while extracellular diffusivity (*D*_*e*_) runs the other way (median *D*_*e*_ 0.94 inner versus 1.12 µm^2^/ms outer) and exchange is slower at the centre (median *t*_*ex*_ 22.0 versus 16.1 ms). A core richer in processes and poorer in extracellular space, surrounded by a looser rim where progenitors are expected to project, is consistent with the radial cytoarchitecture the directed protocol is designed to produce. These estimates describe a clear radial difference in the effective compartmental composition of Batch 1.

Batch 2 shows no such organization. High-FA and high-MK regions are distributed in patches across each organoid rather than as a centroid-to-periphery gradient, and the between-organoid spread is visibly larger (Figure 3). Median *f*_*n*_ is above Batch 1 overall (0.58), but it does not resolve into a shared spatial pattern. Median *t*_*ex*_ is longer than in Batch 1 (39.3 versus 17.3 ms); given the railing described before, we read this as exchange being slower than the Batch 1 regime rather than as a calibrated value.

DAPI cell nuclei staining supports two readings of this heterogeneity. First, the three organoids differ from one another in the spatial arrangement of their nuclei (Figure 2), so the inter-organoid variability in the parameter maps is consistent with genuine tissue differences contributing to the observed inter-organoid variability. Second, it provides anatomical support for a cavity-like interpretation of the localized bright regions in the MD maps of Organoids 1 and 3 (Red arrow in Figure 2). These voxels carry a strong *b* = 0 signal that falls away sharply by *b* = 1000 s/mm^2^ and return elevated MD, *D*_*i*_ and *D*_*e*_, and low FA and *f*_*n*_, and the corresponding DAPI projections show them to be largely devoid of nuclei. A fast-decaying, high-diffusivity signal over an area with few cell bodies indicates fluid-filled or cell-poor cavities rather than tissue, although DAPI alone cannot distinguish lumen-like, cystic, and necrotic cavities.

**Figure 2.**
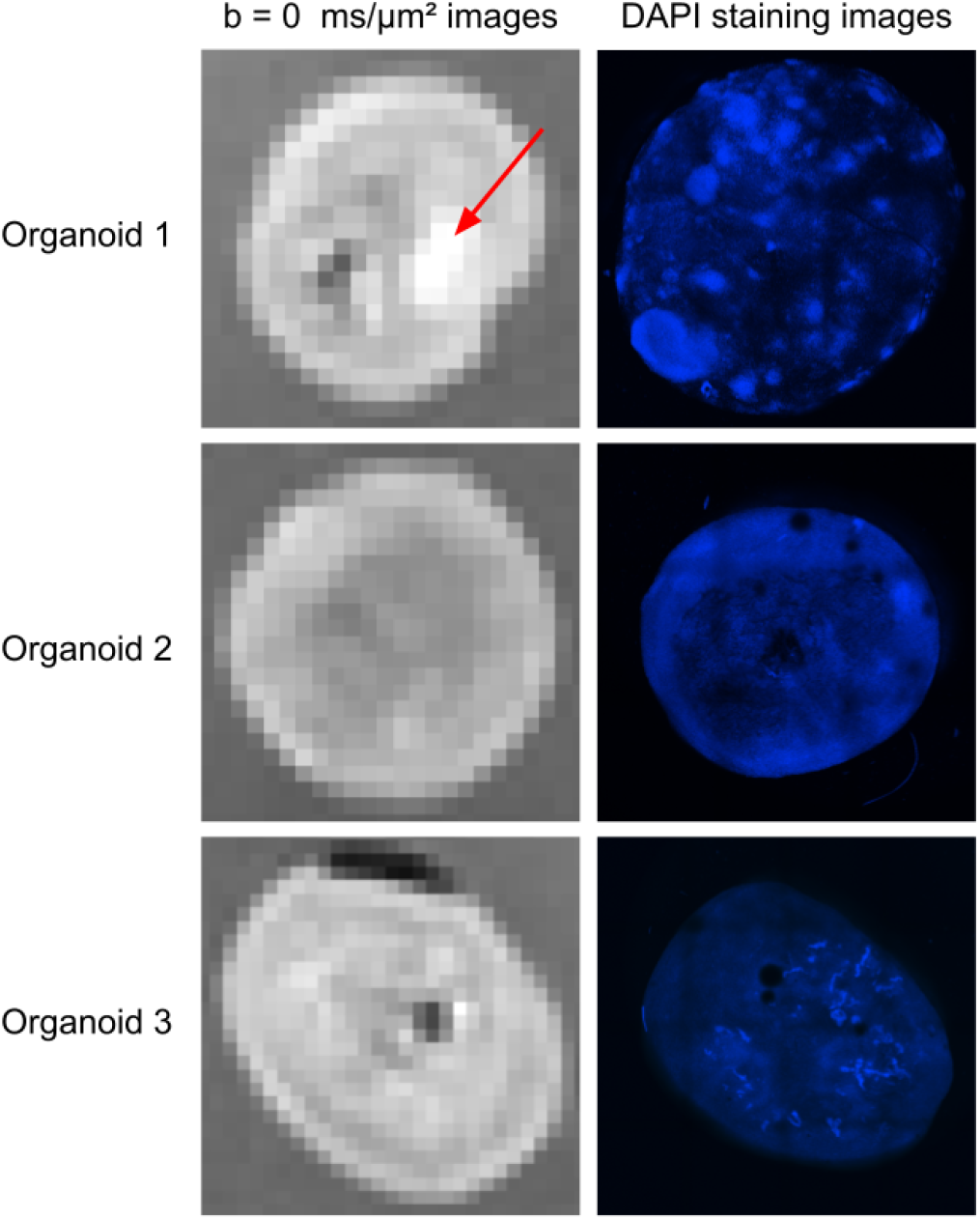
Batch 2: Organoids and their corresponding DAPI staining. Column 1 shows the *b* = 0 ms/*µ*m^2^ images. Column 2 shows the corresponding maximum-intensity Z-plane 2D projections of the DAPI staining, manually oriented to match the orientation of the b = 0 images. The red arrow in Organoid 1 highlights a localized bright region in the b = 0 map.

**Figure 3.**
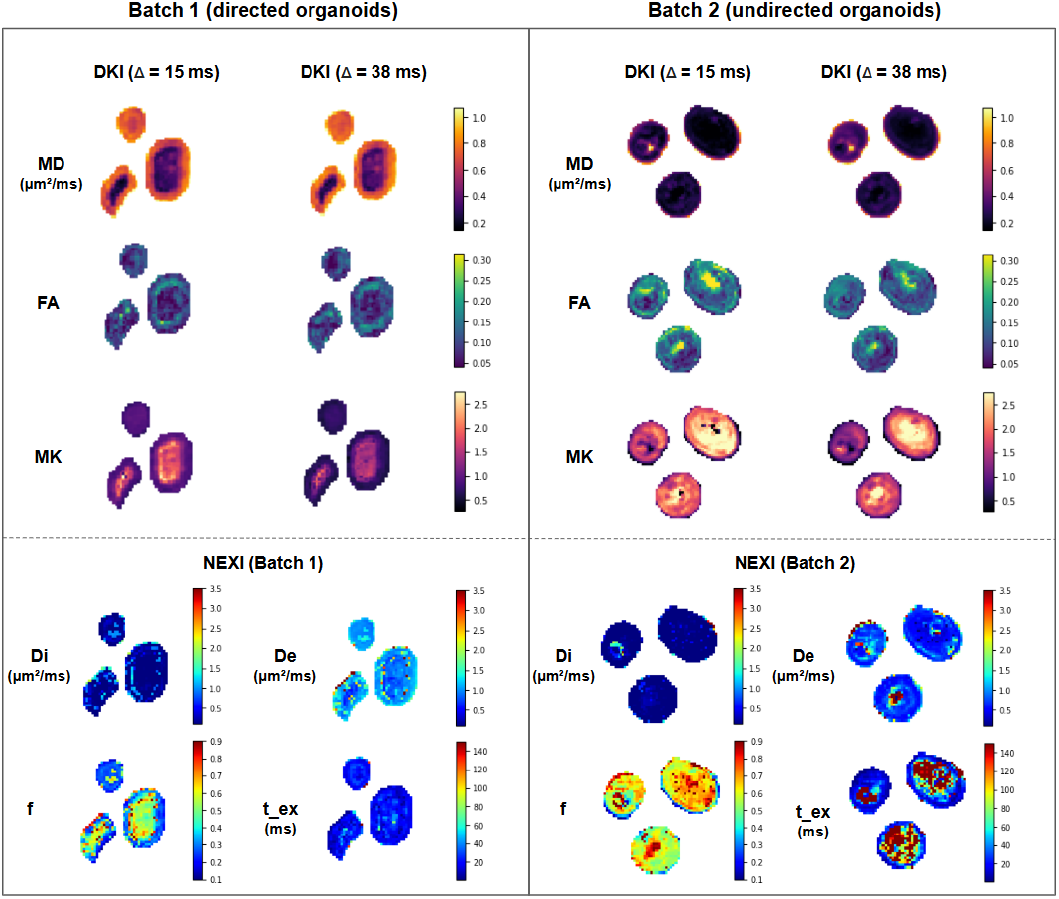
Voxelwise DKI and NEXI parameter maps in Batch 1 (directed) and Batch 2 (undirected) organoids. DKI maps (top) show MD, FA, and MK at Δ= 15 and 38 ms fitted per batch using the corresponding batch-specific b-window; NEXI maps (bottom) show *D*_*i*_, *D*_*e*_, *f*_*n*_, and *t*_*ex*_.

Radial profiling makes the difference quantitative. At Δ = 15 ms the three Batch 1 organoids trace nearly superimposed profiles for MD, FA, and MK along the normalized radius, whereas Batch 2 profiles diverge at most radial bins, with between-organoid differences most evident for MK and FA (Figure 4, Panel B). Lin’s Concordance Correlation Coefficient (CCC) between organoid pairs confirms this at every diffusion time and for every metric, without exception (Figure 4, Panel C): MD, 0.74–0.77 in Batch 1 against 0.48–0.58 in Batch 2; MK, 0.85–0.88 against 0.31–0.65; FA, 0.57–0.63 against 0.16–0.47. The gap is widest for FA, the metric most directly tied to coherent alignment.

**Figure 4.**
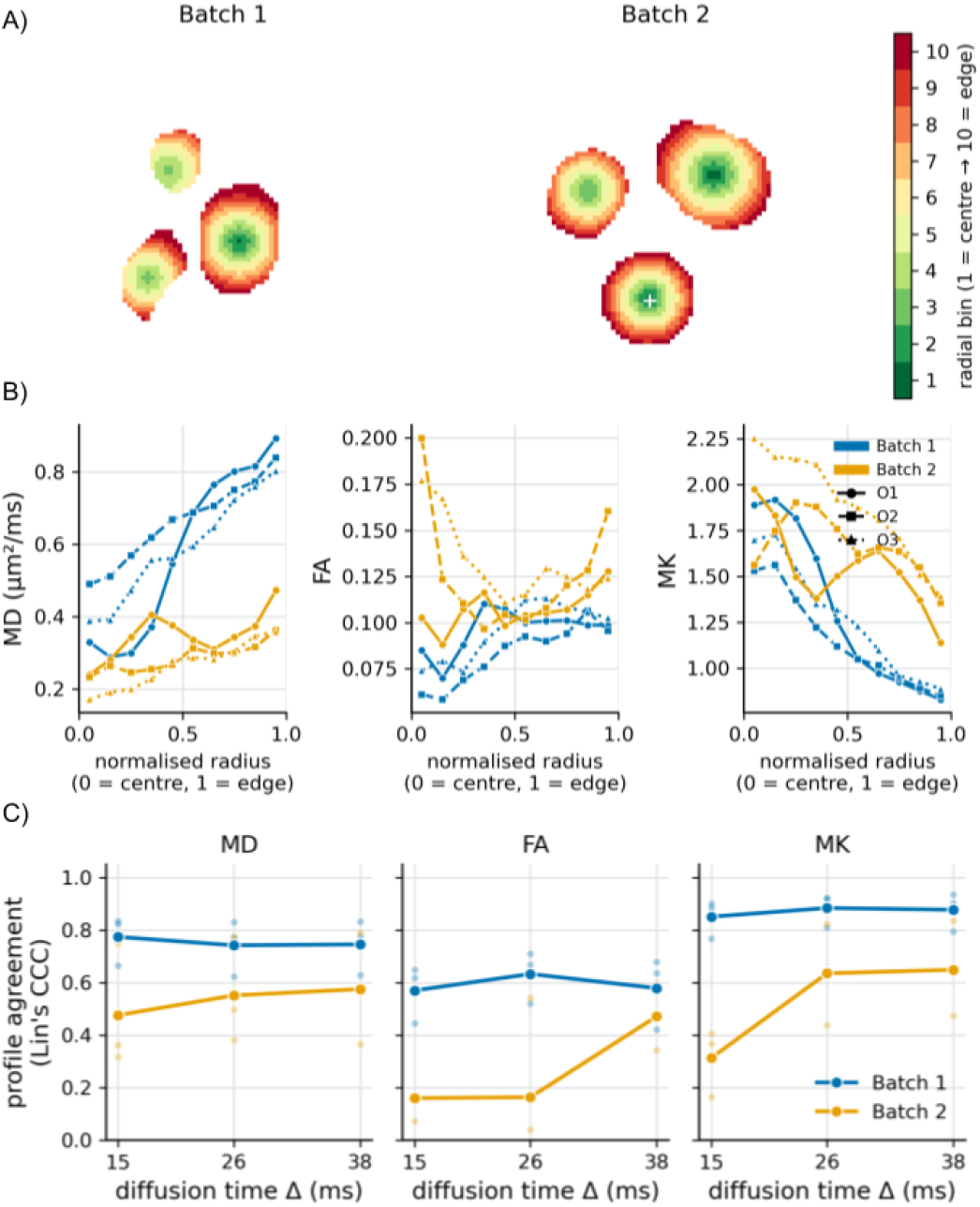
**(A)** Radial bin masks overlaid on the organoid tissue masks for Batch 1 (left) and Batch 2 (right). Each voxel is color-coded according to its radial bin (1 = organoid center, 10 = periphery), derived from the normalized Euclidean distance to the organoid centroid. White crosses indicate the centroid voxel for each organoid. **(B)**: Mean radial profiles of MD (µm^2^/ms), FA, and MK at Δ = 15 ms, plotted as a function of normalized radial distance for each organoid. Blue lines represent Batch 1 organoids (O1: solid circles, O2: dashed squares, O3: dotted triangles) and orange lines represent Batch 2 organoids (same line styles). **(C)** Mean Lin’s concordance correlation coefficient (CCC) of the radial profiles between all intra-batch organoid pairs (O1–O2, O1–O3, O2–O3), shown as a function of diffusion time (Δ = 15, 26, 38 ms) for MD, FA, and MK; faint dots show the three individual pairwise values. Blue and orange lines correspond to Batch 1 (directed) and Batch 2 (undirected) organoids, respectively.

### DKI metrics covary with NEXI-derived compartment parameters

We correlated DKI metrics with NEXI parameters voxelwise across all organoid tissue. Correlations involving the NEXI exchange time *t*_*ex*_ strengthened with Δ. MK–*t*_*ex*_ rises from +0.16 to +0.30 in Batch 1 and from +0.21 to +0.45 in Batch 2 between Δ = 15 and 38 ms (+0.36 to +0.53 pooled), and MD–*t*_*ex*_ becomes correspondingly more negative (−0.15 to −0.40 in Batch 2, −0.37 to −0.50 pooled; the trend is weak in Batch 1, −0.08 to −0.16). MD–*D*_*e*_ also strengthens with Δ (+0.20 to +0.36 in Batch 2, +0.52 to +0.60 pooled). This Δ-dependence is consistent with an increasing contribution of compartment-boundary and exchange-related effects at longer diffusion times.

The compartment-fraction correlations, by contrast, are flat in Δ and consistent across batches: MK correlates positively with *f*_*n*_ (*ρ* = +0.60 and +0.50 in Batch 1 and Batch 2, +0.59 pooled) and negatively with *D*_*e*_ (−0.33 and −0.38), while MD does the reverse (*f*_*n*_: −0.51 and −0.58; *D*_*e*_: +0.51 and +0.29). FA associates weakly with every NEXI parameter (|*ρ*| ≤ 0.31) and reaches neither the strength nor the between-batch consistency of the MD and MK associations.

### As opposed to white matter analyses and organoid studies, average apparent MD increased with diffusion time

Tissue-averaged MD rises with diffusion time in both batches (Figure 5, Panel A), the opposite of the decrease reported in white matter ^30;31^. The effect is not an averaging artefact: voxelwise MD slopes are positive in 90.7% of Batch 1 and 81.5% of Batch 2 tissue voxels, with mean slopes of 2.2 × 10^*−*3^ and 1.9 × 10^*−*3^ µm^2^/ms^2^ and median slopes of 2.1 × 10^*−*3^ and 1.4 × 10^*−*3^ µm^2^/ms^2^ (Figure 5, Panel C) making both slope distributions sit well clear of zero.

**Figure 5.**
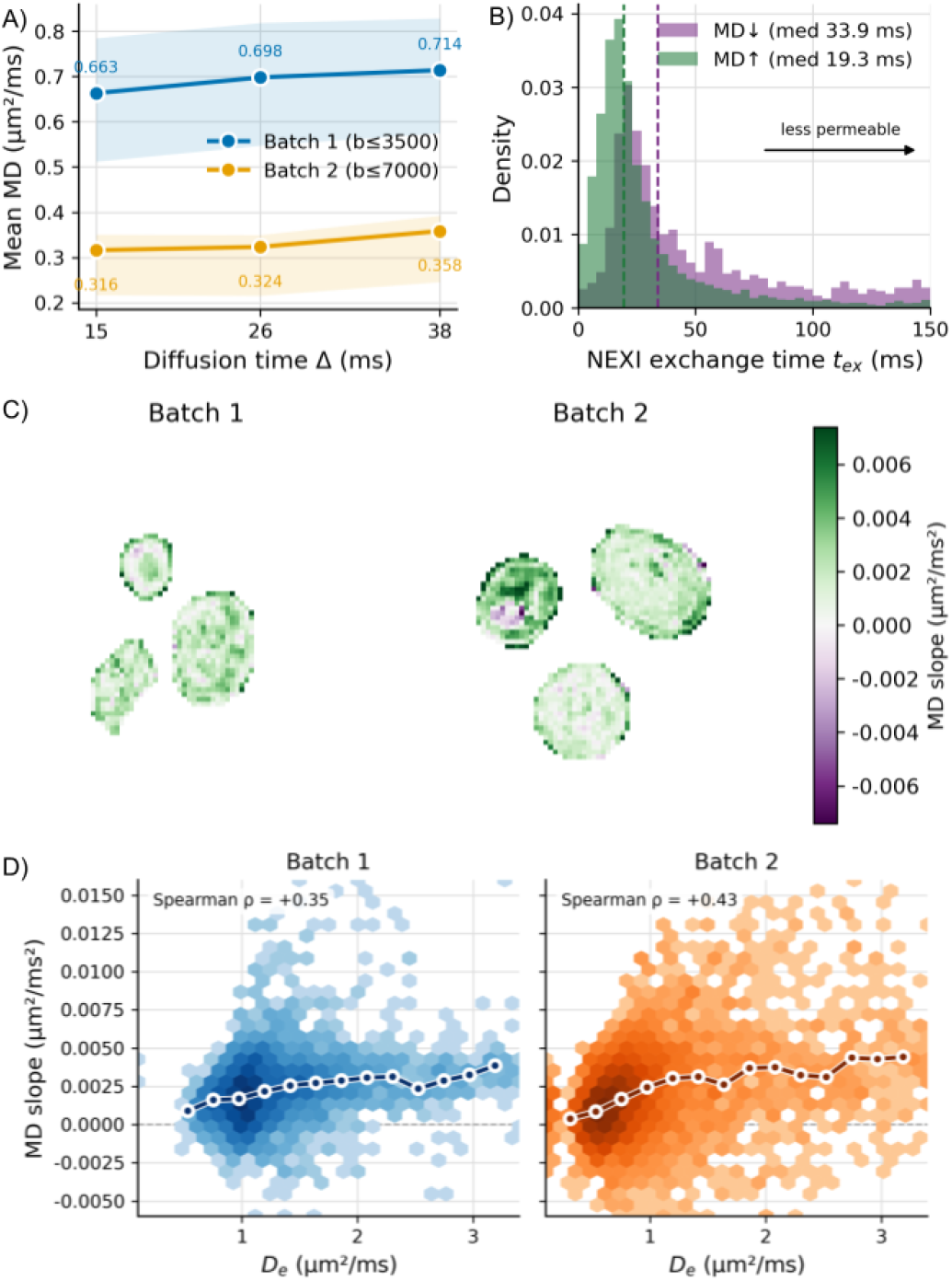
MD evolution study. **(A)** Average MD (µm^2^/ms) estimated with the DKI model as a function of diffusion time Δ in Batch 1 (blue) and Batch 2 (orange), each fitted over its own b-window (*b ≤* 3500 and *b ≤* 7000 s/mm^2^, respectively); shaded regions represent the voxelwise interquartile range. **(B)** Distribution of the NEXI exchange time *t*_*ex*_, pooled over both batches and coloured by the sign of the MD change with Δ: increasing (green) and decreasing (purple), with dashed lines marking the medians and the arrow indicating the direction of slower exchange. Railed *t*_*ex*_ (= 150 ms, the upper bound of the estimable range) was excluded from the distributions and medians: MD*↑ n* = 11,411 (1074 railed), MD*↓ n* = 1218 (978 railed). **(C)** Voxelwise DKI-derived MD-slope maps for one representative slice of Batch 1 (left) and Batch 2 (right); green indicates MD increasing with Δ and purple decreasing (µm^2^/ms^2^). **(D)** MD slope (µm^2^/ms^2^) versus the NEXI extracellular diffusivity *D*_*e*_ (µm^2^/ms) for every tissue voxel, Batch 1 (left, blue) and Batch 2 (right, orange), shown as a hexbin density with the binned-median trend overlaid as a white-outlined line; Spearman *ρ* = +0.35 and +0.43, respectively, and it is computed on all voxels.

As described in Methods 4.2, diffusion times were acquired in opposite orders in the two batches (Batch 1 descending, Δ = 38 → 26 → 15 ms; Batch 2 ascending), so a monotonic drift with time in the scanner would produce MD trends of opposite sign in the two batches; however, both increase. In Batch 1 the last-acquired measurement (Δ = 15 ms) carries the lowest MD, so any upward drift over the session would act against the observed effect, making the measured ΔMD (= MD(Δ=38 ms) − MD(Δ=15 ms)) a lower bound.

The increase is already present in the low-*b* ADC (Supplementary Figure S2). Estimated from *b* = 0 and *b* = 1000 s/mm^2^ alone, the batch-averaged ADC increased from 0.602 to 0.643 µm^2^/ms between Δ = 15 and 38 ms in Batch 1 (+6.8%, 95% CI +6.6 to +7.0) and from 0.312 to 0.329 µm^2^/ms in Batch 2 (mean +5.6%, +5.0 to +6.2), reproducing the direction of the DKI result in both batches. Thus, we also observe a positive diffusion-time dependence using a separate estimator, independent of the DKI fit and the different fitting windows used for the two batches. However, this measurement may remain sensitive to non-Gaussian signal contributions and does not necessarily reflect an increase in the underlying second-cumulant diffusivity.

Exchange times from NEXI separate the two populations. Among voxels whose MD decreases with Δ, 44.5% (978/2196) rail at the 150 ms upper bound, against 8.6% (1074/12,485) of increasing voxels; the same asymmetry is present within each batch (Batch 1: 1.0% versus 0.2%; Batch 2: 58.1% versus 14.5%). In Figure 5 Panel B, after exclusion of railed *t*_*ex*_ values, increasing voxels show a median *t*_*ex*_ of 19.3 ms (IQR 13.0–31.2 ms) while decreasing voxels show a broader, right-shifted distribution (median 33.9 ms, IQR 21.6–65.6 ms). The association holds within each batch (Spearman *ρ* = −0.34 in Batch 1, −0.62 in Batch 2, −0.55 pooled). MD slope also correlates positively with *D*_*e*_ (+0.35 and +0.43; +0.44 pooled) as shown in Figure 5 Panel D, confirming the previous increase in correlation between MD and *D*_*e*_ that was increasing with Δ. In contrast, the MD slope was weakly and inconsistently correlated with *f*_*n*_ (+0.30 and +0.15), and not interpretable with *D*_*i*_.

The previous results associate increasing MD with shorter NEXI exchange times and higher estimated extracellular diffusivity. We therefore used controlled simulations to test whether these conditions can generate a positive ΔMD within a two-compartment exchange model.

### Exchange simulations reproduce increasing apparent diffusivity in analytical and explicit cellular geometries

We simulated substrates covering intracellular diffusivity *D*_*i*_, extracellular diffusivity *D*_*e*_, intracellular volume fraction *f* , membrane permeability *κ*, and spherical compartment radius *R*.

Within the simulated parameter space, non-zero permeability was necessary to reproduce a positive ΔMD. None of the 2058 impermeable substrates (*κ* = 0) produced a positive ΔMD in either *b*-window. Beyond that, the sign of ΔMD is set by *D*_*e*_ relative to the apparent intra-compartment diffusivity *D*_*i*,app_ rather than by the nominal diffusivities (Figure 6, Panel B): no substrate with *D*_*e*_ *< D*_*i*,app_ increased, the increasing fraction grew monotonically with *D*_*e*_*/D*_*i*,app_ to 75% above a ratio of 10 in the *b* ≤ 7000 window, and 12% of substrates with *D*_*i*_ = *D*_*e*_ still increased because restriction pulls *D*_*i*,app_ below *D*_*i*_ in small spheres. In the (*D*_*e*_, *D*_*i*_) plane, ΔMD is negative on and above the *D*_*i*_ = *D*_*e*_ diagonal and positive only for *D*_*e*_ ≫ *D*_*i*_; the NEXI interquartile ranges of both batches fall inside the positive region (Figure 6, Panel A).

**Figure 6.**
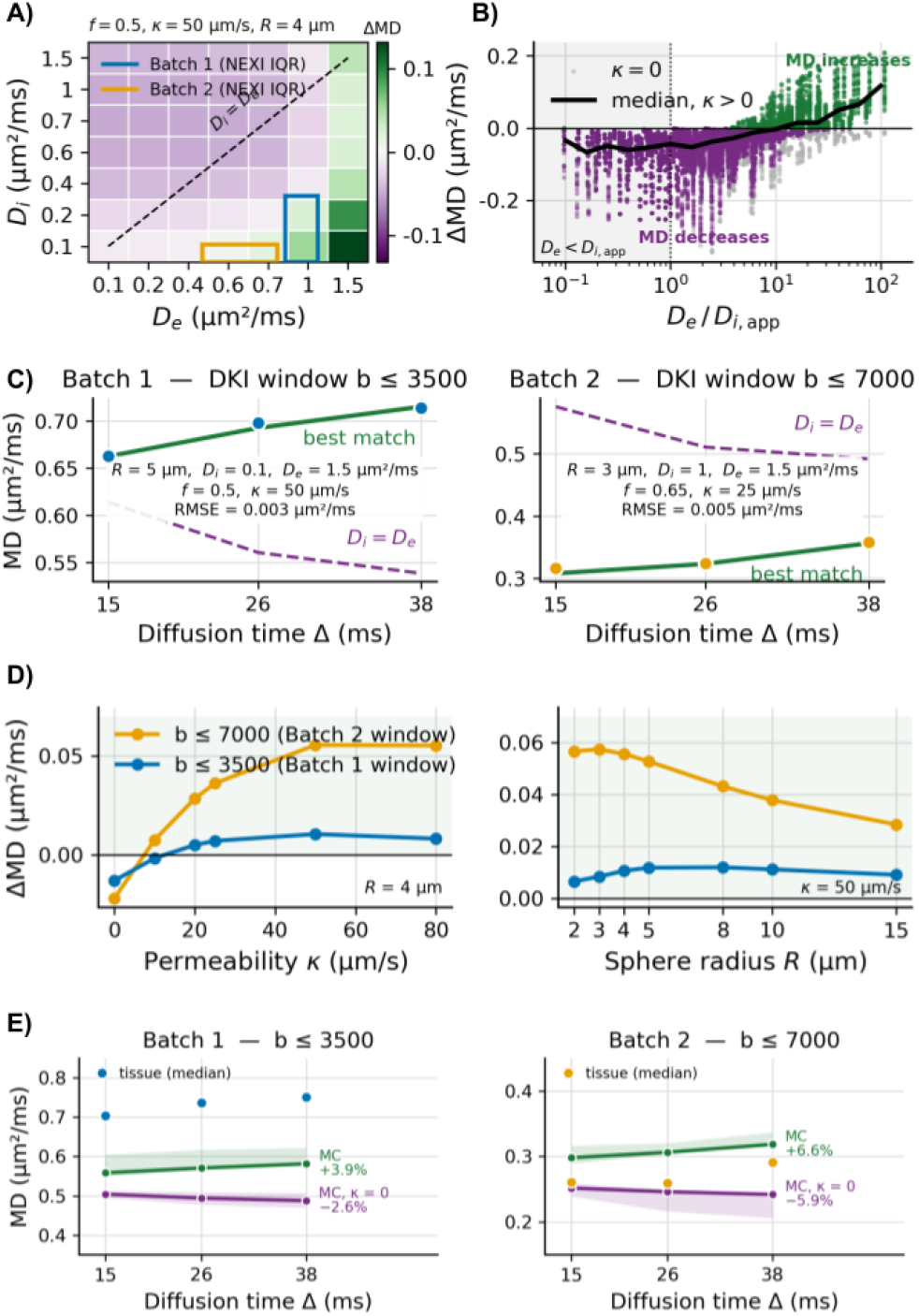
Two-compartment exchange simulations of diffusion-time-dependent MD. Throughout, ΔMD = MD(38, ms) *−* MD(15, ms) from full-protocol DKI fits; green/purple indicate increases/decreases. **(A)** ΔMD across the (*D*_*e*_, *D*_*i*_) plane (*f* = 0.5, *κ* = 50 µm/s, *R* = 4 µm; *b ≤* 7000). The dashed line marks *D*_*i*_ = *D*_*e*_; rectangles show NEXI interquartile ranges for Batch 1 (blue) and Batch 2 (orange). **(B)** ΔMD of all 12,348 substrates versus *D*_*e*_*/D*_*i*,app_, with *D*_*i*,app_ the apparent diffusivity of the isolated restricted sphere. Grey points denote impermeable substrates (*κ* = 0), the black line denotes the median for permeable substrates, and the shading indicates the region *D*_*e*_ *< D*_*i*,app_, where MD does not increase. **(C)** Measured tissue-averaged DKI MD (blue/orange points for Batch 1/2), best-matching substrates (green; parameters and RMSE shown), and a common reference substrate (purple dashed; *R* = 5 µm, *D*_*i*_ = *D*_*e*_ = 1 µm^2^/ms, *f* = 0.5, *κ* = 50 µm/s), fitted within each batch’s DKI window. **(D)** ΔMD in the organoid regime (*D*_*i*_ = 0.1, *D*_*e*_ = 1.0 µm^2^/ms, *f* = 0.5) versus permeability (*R* = 4 µm; left) and radius (*κ* = 50 µm/s; right), using both DKI windows on the same substrate (blue: *b ≤* 3500, Batch 1; orange: *b ≤* 7000, Batch 2). **(E)** Tissue median DKI MD (blue/orange points for Batch 1/2) versus MC simulations (green) and their impermeable counterparts (*κ* = 0, purple). Curves show the median of each batch’s ten best-matching analytical substrates, with interquartile shading, fitted within the respective DKI windows. Percentages indicate median simulated MD changes from Δ = 15 to 38 ms.

The best-matching analytical substrates reproduced the measured MD curves quantitatively (see Figure 6, Panel C). The fit window itself scales the effect: across all 12,348 substrates, 16% increased in the *b* ≤ 7000 window against 9% in the *b* ≤ 3500 window, and among the 1,992 substrates that increased in at least one window, ΔMD was larger in the wider window in every case. Analyzed in its own DKI window, the best-matching substrate for Batch 1 (*R* = 5 µm, *D*_*i*_ = 0.1, *D*_*e*_ = 1.5 µm^2^/ms, *f* = 0.5, *κ* = 50 µm/s) gave MD = 0.663, 0.693, 0.715 µm^2^/ms against measured 0.663, 0.698, 0.714 (RMSE 0.003), and for Batch 2 (*R* = 3 µm, *D*_*i*_ = 1.0, *D*_*e*_ = 1.5, *f* = 0.65, *κ* = 25 µm/s) 0.308, 0.323, 0.356 against 0.316, 0.324, 0.358 (RMSE 0.005). Both reproduce the measured curve to within one to two standard errors of the tissue mean (SEM 0.003 and 0.002 µm^2^/ms), and recovered ΔMD to within 0.002 and 0.007 µm^2^/ms of the measured value. Although the two best matches differ tenfold in nominal *D*_*i*_, their apparent intracellular diffusivities were nearly identical (0.077 and 0.072 µm^2^/ms), giving *D*_*e*_*/D*_*i*,app_ of 19.5 and 20.8. The ten best-matching substrates for each batch all had *D*_*e*_ *> D*_*i*,app_ and non-zero permeability; among the fifty best, *D*_*e*_ *> D*_*i*,app_ and *κ >* 0 held without exception. The *D*_*i*_ = *D*_*e*_ control decreased in both windows (ΔMD −0.075 and −0.084).

The magnitude scales with permeability as expected of an exchange effect. In the organoid regime (*D*_*i*_ = 0.1, *D*_*e*_ = 1.0 µm^2^/ms, *f* = 0.5, *R* = 4 µm), ΔMD moves from −0.022 µm^2^/ms at *κ* = 0 through +0.008, +0.029 and +0.056 at *κ* = 10, 20 and 50 µm/s, saturating by *κ* = 80 (Figure 6, Panel D). At *κ* = 50 µm/s, the increase holds at every radius tested, weakening from +0.057 at *R* = 2 µm to +0.028 at *R* = 15 µm. The *b* ≤ 3500 window gives the same trend with smaller magnitudes.

To assess whether the positive diffusion-time dependence predicted by the analytical model persists in an explicit cellular geometry, we simulated the ten best-matching analytical substrates for each batch using Monte-Carlo (MC) random walks in permeable sphere packings (Figure 6, Panel E; Supplementary Table S1). DKI-derived MD increased between Δ = 15 and 38 ms in all twenty substrates, with median relative changes of +3.9% (range +1.3 to +5.5%) for Batch 1 and +6.6% (+4.4 to +11.8%) for Batch 2. This increase required exchange: when the membranes of the same substrates were made impermeable (*κ* = 0), MD decreased with Δ in all twenty, with median relative changes of −2.6% (range −6.4 to −1.2%) for Batch 1 and −5.9% (−24.6 to −2.0%) for Batch 2. However, the simulations did not reproduce the absolute diffusivity. At Δ = 15 ms, Batch 1 substrates underestimated the median tissue MD by 17–25% when *D*_*i*_ = 0.1 µm^2^/ms and by 7–13% when *D*_*i*_ ≥ 1 µm^2^/ms, whereas Batch 2 substrates overestimated it by 7–27%. The low-*b* ADC (*b* = 1000 s/mm^2^) reproduced the trend in Batch 1 (simulations: median +4.5%; tissue: mean +6.8%, median +6.8%), whereas in Batch 2 it barely changed in the simulations (median +0.2%, mean +0.3%) but increased in the tissue mean (+5.6%) much more than in the tissue median (+1.8%), suggesting that the larger increase in the mean was driven by a subset of voxels with particularly strong positive changes.

To determine whether the simulated increase in DKI-derived MD also reflected an increase in the underlying physical diffusivity, we further analyzed the Monte Carlo trajectories of the best-matching Batch 1 substrate (Supplementary 7.1). Despite an increase in DKI-derived MD of 5.4% between Δ = 15 and 38 ms, the displacement-derived diffusivity (*D*_MSD_) decreased by 1.2%, while the second-cumulant diffusivity (*D*^(2)^), calculated independently from the accumulated PGSE phase covariance, decreased by 0.7%. Over the same diffusion-time range, the small-*b* kurtosis decreased from 2.05 to 1.52, whereas the finite-*b* ADC at *b* = 1000 s/mm^2^ increased by 5.2%. As the DKI fitting window was progressively reduced towards *b* → 0, the positive MD trend weakened and eventually reversed. In the matched impermeable substrate (*κ* = 0), both the physical and finite-*b* apparent diffusivities decreased with diffusion time. These results show that an increase in finite-*b* DKI-derived MD can coexist with a decrease in the underlying displacement-derived and second-cumulant diffusivities within the investigated permeable cellular geometry.

## Discussion

In this study, we investigated the potential of cortical organoids as experimental platforms for characterizing early cortical microstructure with diffusion MRI and evaluating biophysical diffusion models. Diffusion MRI of fixed cortical organoids yielded two findings. First, the radial organization of the diffusion signal was more reproducible across the three organoids grown under the directed protocol than across the three undirected organoids, consistent with the greater reproducibility previously reported for directed differentiation strategies. Second, finite-*b* DKI-derived MD increased with diffusion time in both batches, opposite to the decrease reported in previous organoid measurements ^28;29^. Analytical exchange simulations and explicit-geometry Monte-Carlo simulations showed that a positive MD slope can arise when water exchanges across permeable membranes between restricted intracellular and relatively faster extracellular environments. Further trajectory analysis showed that an increase in fitted MD can coexist with decreasing displacement-derived and second-cumulant diffusivities, highlighting the importance of non-Gaussian signal contributions when interpreting diffusion-time dependence. Together, these results show that the organoid tissue studied in this project shows a particular regime of GM diffusion, including strong intracellular restriction and potentially appreciable water exchange where current biophysical models are at the limit of their assumptions.

Both young and mature living cortical organoids ^28^ and fixed forebrain organoids ^29^ have shown higher diffusivity at the periphery than in the core, with histology matching this to a necrotic core surrounded by viable cells. Organoids of the size scanned here commonly develop such a core (Methods 4.1), so the inner-outer contrast in Batch 1 likely reflects, at least in part, this core-rim gradient where estimated MD is higher in the periphery of the organoid than at its core. However, the NEXI-derived exchange-time pattern is not consistent with a purely necrotic interpretation. Wu et al. found faster exchange in necrotic cores, consistent with disrupted membranes ^29^, whereas our Batch 1 inner ring exchanges more slowly than the outer ring (median *t*_*ex*_ 22.0 versus 16.1 ms). Because slower exchange can also arise from larger restricting compartments at unchanged permeability, and no histology is available for Batch 1, these data indicate a core that is not cell-depleted but cannot establish whether it is viable or in an early necrotic state.

An important difference between the two batches is how consistently this spatial organization repeats from one organoid to the next. Batch 1 followed a forebrain spheroid protocol ^32^, whereas Batch 2 followed an undirected protocol that yields a more variable organization of cellular foci displaying diverse cell density and organization ^33^. To our knowledge, this is the first study to use dMRI to directly compare human brain cortical organoids generated using guided and unguided differentiation protocols. In this study, the radial concordance tracks this difference for every metric at every diffusion time, and nuclear staining shows that the unguided organoids differ in the spatial arrangement of their nuclei, so the disagreement between their diffusion maps supports biological differences contributing to the observed variability. These observations show that dMRI is sensitive to differences in the spatial organization of organoid tissue and can quantify inter-organoid reproducibility. In the present dataset, the observed contrast is associated with batches generated using different differentiation strategies and is consistent with previously reported protocol-related differences in organoid reproducibility ^34^.

Within these organoids, MD and MK showed stronger associations with NEXI-derived compartment parameters than FA did. FA remained low and correlated weakly with the NEXI parameters, whereas MD and MK covaried with the effective restricted-compartment fraction *f*_*n*_ in opposite directions. In the Batch 1 inner ring, a high *f*_*n*_ aligned with low FA, which in NEXI’s terms describes densely packed restricted compartments without a common orientation. In tissue composed mainly of neurons, progenitors, and few glia (Methods 4.1), NEXI’s neurite compartment might be better read as a restricted compartment than as neurites in the anatomical sense. In fixed organoids, dense rosette cores likewise appear isotropic ^25^. Because DKI and NEXI are fitted to the same signal, their static association is partly structural. Thus, the informative feature is its dependence on diffusion time. Correlations with *f*_*n*_ remained stable across Δ, whereas correlations with the exchange time increased monotonically (Results 2.2). This trend is consistent with the increasing sensitivity of diffusion metrics to compartment boundaries and exchange as Δ increases, providing water molecules with more time to encounter cellular barriers and to exhibit stronger signatures of restricted intracellular diffusion in the measured MK in less permeable tissues. Agreement between an empirical representation and a biophysical model on this Δ-dependence is consistent with a contrast governed by compartmentalization and exchange.

The small bright regions visible in the MD maps of Batch 2 Organoids 1 and 3 (Figure 3, top) are characterized by high MD, low FA, low MK, and low *f*_*n*_, yielding a signature consistent with a fluid-rich or weakly restricted compartment. These regions are surrounded by tissue with higher *f*_*n*_ and anisotropy. Several structures could plausibly give rise to this pattern. Fluid-filled cysts are a recognized outcome of cerebral organoid differentiation ^24;33^, and cystic regions have been shown to exhibit higher diffusivity than compact organoid tissue on DTI ^24^. Another option is that undirected cerebral organoids commonly develop multiple rosette-like neuroepithelial structures, in which neural progenitor cells are radially organized around central lumen-like spaces ^14;35^. Consistent with this interpretation, ventricular-zone-like structures in cortical organoids are characterized by polarized neuroepithelium arranged around a central lumen ^23^. In living cortical organoids, regions surrounding ventricular-like zones likewise show elevated macroscopic anisotropy and mean diffusivity, with the anisotropic signal interpreted as reflecting radially aligned tissue. They confirmed this interpretation with subsequent light-sheet imaging and identified the corresponding ventricular-like zones using ZO-1 staining ^28^. A third possibility is necrotic cavity formation, since both batches could develop necrotic cores (Methods 4.1). The hollow structures visible in the DAPI images (Figure 2) of all three Batch 2 organoids support the presence of lumen- or cavity-like structures, but DAPI establishes only the absence of nuclei and cannot distinguish a fluid-filled lumen lined by progenitors from a necrotic cavity. The more prominent signature in Organoids 1 and 3, together with the weaker signature in Organoid 2, could reflect differences in cavity size or spatial dispersion relative to the 200-µm voxel size, with smaller or more dispersed cavities producing partial-volume averaging between fluid and surrounding tissue. Biophysical models that explicitly include a free-water compartment could provide greater specificity for distinguishing these possibilities in future studies.

All organoids were fixed in 4% paraformaldehyde for up to two hours, with a fixation time adjusted to organoid size, and scanned at approximately 20 ^*°*^C (Methods 4). These experimental conditions should be considered when interpreting the absolute diffusivities, as both chemical fixation and temperature alter tissue diffusion properties ^36^. However, they do not provide a direct explanation for the positive diffusion-time dependence observed here. Our DKI-derived apparent MD increased with diffusion time, in contrast to previous measurements in fixed forebrain organoids, where diffusivity decreased over t = 7–200 ms, with most of the time dependence occurring below approximately 20 ms ^29^. In living cortical organoids, ADC also decreased with increasing diffusion time between 2.7 and 25 ms, except for a transient increase at the periphery of a young organoid, which the authors suggested could reflect kurtosis-related bias in a mono-exponential ADC fit ^28^. The use of DKI in our study accounts for the leading quadratic non-Gaussian contribution ^37^ that biases mono-exponential ADC estimates. However, higher-order signal terms may still influence the fitted diffusivity over a finite *b*- range, as supported by the Monte Carlo analysis and the dependence of DKI-derived MD on the fitting window (Supplementary 7.1). Importantly, even relatively low *b*-values do not guarantee that mono-exponential diffusivity estimates closely approximate the underlying second cumulant when non-Gaussian signal contributions remain appreciable. The opposite Δ acquisition order used for the two batches also argues against a systematic monotonic heating effect as the source of the observed trend (Methods 4.2). A closer precedent is provided by fixed postmortem human cortex, in which diffusivity showed weak-to-absent diffusion-time dependence in the frontal lobe but an increase over 12–100 ms in the temporal lobe; the latter was tentatively attributed to restriction domains becoming permeable at longer diffusion times ^38^. Although that interpretation remains speculative, it illustrates that positive diffusion-time dependence can occur in fixed gray matter.

The different sign of the MD(Δ) dependence across studies likely reflects a balance of several competing effects rather than the presence or absence of exchange alone. Restriction, compartmental diffusivities, membrane permeability, and the relative signal contribution of the intra- and extracellular spaces can each influence the measured slope and may act in opposite directions. Most of the diffusivity decrease reported by Wu et al. occurred below approximately 20 ms and was weak at longer diffusion times ^29^, whereas Nikolaeva et al. sampled living organoids over 2.7–25 ms ^28^; our measurements span Δ = 15–38 ms and may therefore probe a later regime in which the early restriction-driven decrease is less dominant. Differences in organoid maturation and microstructure, including the relative contributions of somata, short cellular processes, extracellular space, and membrane permeability, could further shift this balance. In particular, exchange between restricted intracellular and relatively faster extracellular environments can modify the non-Gaussian diffusion signal, potentially producing an increase in finite-*b* apparent MD without requiring the underlying physical diffusivity to increase. The measured diffusion-time dependence may therefore reflect not only differences in cellular geometry, compartmental diffusivities, and permeability, but also their influence on the higher-order signal contributions captured by the fitting. Experimental differences in fixation, temperature, *b*-value range, and echo time may additionally alter absolute diffusivities and the relative signal weighting of the compartments. We therefore interpret the positive apparent MD(Δ) slope as an emergent feature of this combined microstructural and experimental regime, rather than as a unique potential signature of exchange or an increase in the underlying physical diffusivity.

The NEXI fits provide a second indication that the organoid microstructure differs from the tissue for which the model was originally developed. The intra-neurite diffusivity *D*_*i*_ reached its lower fit bound in most voxels, whereas *D*_*e*_ was consistently above this bound.

Intra-neurite diffusivity being estimated systematically lower than the extracellular one has previously been observed with NEXI in fixed postmortem human gray matter, in contrast to the usual *D*_*i*_ *> D*_*e*_ ordering reported in vivo and ex vivo in rat ^38^. In the present organoids, however, substantial signal from somata and short processes can make the stick-based NEXI *D*_*i*_ an effective parameter rather than an intrinsic intracellular diffusivity, so the inversion should be interpreted primarily as evidence of model mismatch or strong intracellular restriction. Hertanu et al. proposed that global ischemia following death may induce neurite beading and thereby reduce the fitted *D*_*i*_ ^38^. This explanation is also unlikely to apply directly to organoids, which do not undergo an agonal period, and the rat ex vivo results indicate that ex vivo preparation alone does not necessarily produce the inversion.

One possible explanation is model mismatch: in cortical organoids, the intracellular signal may arise substantially from somata and short cellular processes, whereas NEXI represents this compartment using zero-radius sticks and does not explicitly account for restricted diffusion within cell bodies ^3^. Over the investigated diffusion times and *b*-values, the model may partially absorb these restriction effects into a low effective *D*_*i*_. Additional intracellular barriers could further contribute to restricted diffusion, although their influence cannot be determined from the present data. Chemical fixation may further modify both diffusion and membrane permeability in a compartment-dependent manner ^36^, while the approximately 20 ^*°*^C acquisition temperature reduces diffusivity relative to physiological temperature ^39^; neither effect, however, provides a straightforward mechanistic explanation for the observed *D*_*i*_ *< D*_*e*_ inversion. Because *D*_*i*_ reached the lower bound in a substantial fraction of voxels, we therefore interpret this parameter conservatively as evidence for low apparent intracellular diffusivity, rather than as a calibrated estimate of intrinsic neurite diffusivity.

For Batch 1, these parameters suggest that the organoids may lie outside the fully coarse-grained regime assumed by the Kärger exchange model underlying NEXI. For its best-matching substrate, *κR/D*_*i*_ is approximately 2.5; the corresponding mean intracellular residence time is ∼33 ms, compared with a characteristic intracompartmental mixing time of ∼58 ms for a reflecting sphere. Thus, intracellular water may exchange before fully sampling the compartment, and the mixing time also exceeds our Δ = 15– 38 ms range. This holds across the substrates that fit Batch 1 equally well (*κR/D*_*i*_ 2.5–8.0, mixing times 58–231 ms). For Batch 2, the data do not resolve the question: substrates fit within one standard error of the optimum span *κR/D*_*i*_ from 0.03 to 2.0 and mixing times from 1.3 to 148 ms, so both regimes remain compatible with the measured curve. Importantly, *D*_*i*_ and MD are estimated using different models, although both are fitted to the same diffusion signal; because the two models also assign different geometries to their intracellular compartments, their *D*_*i*_ are not microscopic equivalents. The isolated-sphere apparent intracellular diffusivities were low for the best-matching analytical substrates (0.077 and 0.072 µm^2^/ms). These are substrate-dependent estimates rather than direct measurements in the organoids, but they place the selected substrates in a regime of strong intracellular restriction over the investigated diffusion-time window. Exchange between restricted intracellular and relatively faster extracellular environments provides a plausible mechanism contributing to the observed increase in finite-*b* apparent MD, although the fitted trend does not directly describe the diffusion-time dependence of the underlying physical diffusivity. We therefore interpret the NEXI parameters as effective, model-dependent quantities rather than direct microscopic measurements ^31;40^.

Our analytical simulations rely on Kärger coupling between compartment signals, so their ability to reproduce an MD increase does not establish whether the effect persists when compartment geometry and membrane crossings are treated explicitly. In contrast, the Monte-Carlo substrates explicitly simulate diffusion in permeable spherical compartments and provide an independent test of the mechanism. All twenty substrates selected from the analytical best matches retained a positive diffusion time dependence of DKI-derived MD in permeable sphere packings, while the trend reversed in their matched impermeable counterparts. The reversal of this trend in the matched impermeable substrates indicates that membrane exchange is a key determinant of the positive finite-*b* MD slope in the simulated geometries, although this does not establish exchange as its unique possible origin in biological tissue. Also, three substrates showed increasing MD despite equal intrinsic intra- and extracellular diffusivities (*D*_*i*_ = *D*_*e*_), consistent with the importance of the contrast between *D*_*e*_ and the apparent intracellular diffusivity *D*_*i*,app_, rather than the intrinsic diffusivity alone.

However, the agreement between analytical and MC simulations was qualitative rather than quantitative. For identical substrate parameters, the MC simulations reproduced a median of 55% of the relative MD increase predicted analytically and generally underestimated the measured increase. Substrates with the lowest *D*_*i*_ in Batch 1 showed the largest positive slopes but underestimated absolute MD, whereas substrates with higher *D*_*i*_ came closer to their measured level while showing smaller slopes. In Batch 2, the simulated MD levels were comparatively close to the measurements, but the low-*b* ADC changed little with diffusion time in the simulations (median +0.2%, mean +0.3%), while in the tissue its mean increased by +5.6% and its median by only +1.8%. Therefore, reproducing the experimental MD curve with an analytical model over a given *b*-window does not imply that the corresponding microstructural parameters also reproduce the full diffusion signal in an explicit geometry.

In Batch 2, the simulated MD levels were comparatively close to the measurements, but the low-*b* ADC changed little with diffusion time in the simulations (median +0.2%, mean +0.3%), while in the tissue its mean increased by +5.6% and its median by only +1.8%.

Analysis of the Monte Carlo trajectories further clarified the physical interpretation of this apparent increase. For the best-matching Batch 1 substrate, the displacement-derived diffusivity and the PGSE second-cumulant diffusivity decreased with diffusion time, despite the positive DKI-derived MD slope. This behaviour persisted after equilibration and reversed in the matched impermeable control, supporting an exchange-sensitive origin within the simulated geometry. The decrease in small-*b* kurtosis with diffusion time is consistent with exchange reducing diffusional heterogeneity, which can increase mono-exponential ADC at finite *b*. For the DKI case that already accounts for the quadratic kurtosis term, we think that the increasing fitted MD instead reflects the influence of higher-order signal contributions beyond the truncated DKI expansion. The dependence on the fitting *b*-window, including the recovery of a negative MD trend at sufficiently low *b*, supports this interpretation (Supplementary Section 7.1). Although these simulations demonstrate a plausible mechanism for the observed increase in apparent MD, they do not establish that the same mechanism necessarily explains our findings in organoids.

The differences between analytical and Monte-Carlo simulations with the real tissue signal reflect properties and tissue complexity that are simplified or absent in these two representations. In the MC simulations, cell surfaces hinder extracellular diffusion, whereas the analytical model treats extracellular diffusion as Gaussian and uses coarse-grained exchange between compartments. However, the sphere packings themselves do not capture the cellular processes and populations, rosette-like structures, or the spatial heterogeneity of the organoid tissue. Such features may affect both the absolute diffusivity and its dependence on diffusion time and b-value. Kurtosis was already decreasing at the shortest diffusion time sampled in our data, while fixed forebrain organoids previously exhibited a kurtosis peak near 20 ms, whose timing was found to be more sensitive to thin processes than to sphere size ^29^. More realistic, histology-informed substrates are therefore needed to reproduce the measured MD level, low-b behaviour, and diffusion-time dependence simultaneously.

Several aspects of the present study motivate further work. The comparison was based on three organoids per protocol from a single differentiation, with the protocol also confounded by age at scanning and slice coverage. Future experiments with independent differentiations, larger numbers of organoids, and matched developmental stages would further help separate protocol effects from batch-to-batch and developmental variability. Organoid deformation during embedding, transport, and staining is expected, but it complicates direct spatial matching between MRI and microscopy. Future work could account for it through deformable 3D co-registration, complemented by additional lumen, progenitor, and membrane markers on the same specimens ^28^. Nuclear staining in the present study provided only qualitative anatomical support; quantitative MRI–histology matching will be important for testing the interpretation of the fitted microstructural parameters. Finally, extending the diffusion-time range and incorporating more densely sampled low-*b* measurements, ideally with complementary pulsed-gradient and stimulated-echo acquisitions ^29;38^, could help distinguish changes in the second-cumulant diffusivity from higher-order signal contributions, while further constraining the restriction and exchange mechanisms underlying the observed contrast.

These findings support the use of cortical organoids as experimental platforms for investigating diffusion MRI in developing gray matter, with opportunities for complementary histological characterization. The directed batch exhibited more reproducible radial diffusion profiles, while both batches showed increasing DKI-derived MD with diffusion time. Analytical and explicit-geometry simulations reproduced the direction of this apparent increase under permeable conditions, while trajectory analysis demonstrated that it can coexist with decreasing physical diffusivity. The simulations indicate that exchange-dependent higher-order signal contributions can produce an increase in DKI-derived MD even when the underlying second-cumulant diffusivity decreases. Whether this mechanism explains the experimental observations in organoids remains to be established.

## Methods

### Samples and Preparation

Two batches of three brain cortical organoids each were independently cultured for 4 to 6 months at the NeuroNA Human Cellular Neuroscience Platform (HCNP) from the Fondation Campus Biotech Geneva. Selected organoids had reached a suitable size (∼3–5 mm in diameter) for MRI acquisition at the planned resolution. Batch 1 included three dorsal forebrain organoids at 4.5 months of development. They were generated from the CAU hiPSC line (XX, young adult) obtained from Phenocell company, and following Birey et al. protocol ^32^. At this stage of maturation, these organoids are mainly composed of dorsal cortical neurons, progenitors, and a few glial cells. Given the size of organoids, a necrotic core is commonly found at the center, and healthy cells are distributed at the periphery within ∼500 µm.

Batch 2 included three unpatterned cerebral organoids at 6 months of development. They were generated from the same starting hiPSC line, following the protocol of Lancaster et al. ^33^. Unpatterned cerebral organoids develop a greater brain cell diversity, resulting in a more variable organization of cellular foci displaying diverse cell density and organization.

Both batches followed the same sample preparation protocol. Organoids were fixed in 4% paraformaldehyde for up to two hours based on size and a diffusion rate of ∼1 mm/h. Swirling was used to ensure uniform fixation. Organoids were then washed in phosphate-buffered saline (PBS) and embedded in agarose gel blocks to stabilize them during scanning. Upon arrival at the CIBM Pre-Clinical Imaging EPFL Section (Lausanne, Switzerland), agarose blocks containing the organoids were first stabilized within syringe holders filled with Galden. Samples were then degassed for more than two hours prior to the scanning session to remove trapped air bubbles. This protocol was adapted from the tissue preparation protocol for ex-vivo MRI ^41^.

### MRI acquisition

Both batches were scanned using a 9.4T, 31 cm Bruker horizontal-bore MRI system equipped with a B-GA12S HP gradient set (660 mT/m gradient amplitude, 4570 T/m/s slew rate), an 86 mm quadrature volume transmit coil, and a 2×2 array receive-only Rat CryoProbe (Bruker BioSpin). Multi-shell diffusion MRI images were acquired using a Pulsed Gradient Spin Echo-Planar Imaging (PGSE EPI) sequence with b-values of 1000 (12 directions), 2000 (16 dir.), 3500 (24 dir.), 5000 (30 dir.) and 7000 s/mm^2^ (40 dir.), in addition to four b = 0 images and a reverse phase-encoded one. These five shells were acquired at three different diffusion times Δ = 15, 26, and 38 ms. Batch 1 diffusion data were acquired in descending order of Δ (38 → 26 → 15 ms). Conversely, Batch 2 was acquired in increasing order(15 → 26 → 38 ms). This opposite ordering provides a control against monotonic scanner drift, which would be expected to induce diffusion-time trends of opposite sign in the two batches. Other parameters were fixed: *δ* = 4.5 ms, FOV = 16 x 16 *mm*^2^, matrix: 80 x 80, 18 slices and TE/TR = 54/2400 ms. TE was chosen to accommodate the longest diffusion time (38 ms) and kept constant between acquisitions. Batch 1 data were acquired at 0.2 mm in-plane resolution with a 0.2 mm slice thickness and 0.4 mm spacing between slices. Since organoids were placed far apart, more widely spaced slices were sufficient for coverage. For Batch 2, the slice spacing was reduced to match the in-plane resolution (i.e., 0.2×0.2×0.2 mm). The dMRI scan time was 50 min per Δ, and the CryoProbe outer surface temperature was set to 20°C during both scan sessions.

### Staining

DAPI nuclear staining and confocal imaging were performed on Batch 2 organoids. Following MRI acquisition, organoids were removed from the agarose block without tearing or straining and then rinsed in PBS, permeabilized in PBS and Triton 2% for 2 days at 4°C, and incubated in DAPI (250 ng/mL) for 2 h at room temperature on a shaker. Organoids were then washed three times in PBS over 1 h, left in PBS overnight at 4°C, and immersed in RapiClear 1.47 for tissue clarification at room temperature before imaging.

Imaging was performed on a Nikon AxR laser scanning confocal microscope with a 10X Plan Apo glycerol-immersion objective (NA 0.5, WD 5.5). Images were acquired using a galvano scanner in unidirectional mode with 2× line averaging, a pixel dwell time of 1.6 µs, and a pinhole diameter of 10.9 µm. Confocal Z-stacks were acquired with a Z-step size of 33.4 µm (23–39 optical sections depending on the organoid size). The lateral pixel size was 1.73 µm. Due to the large size of the organoids, the accessible imaging depth of the confocal system did not allow acquisition of the full axial extent of each sample; approximately 200 µm of each organoid volume was captured along the Z-axis. Laser power and detector gain were individually adjusted for each organoid to ensure the DAPI signal remained within the detector dynamic range without saturation (405 nm laser power: 2.4–4.5%; detector gain: 44.7–58.8).

### MRI Pre-processing

Both batches underwent the same pre-processing pipeline. As each diffusion time was acquired in a separate scan, all multi-shell multi-diffusion-time data (N=378 volumes, including b0s) were first pooled together. Pre-processing included *Patch2Self* ^42^ for denoising, and Gibbs ringing correction ^43^. Susceptibility-induced distortion correction was performed using FSL’s topup ^44^, followed by eddy current and motion correction using FSL’s eddy ^45^, applied jointly to the full pooled dataset. Manual segmentation was performed to separate the sample tissue from the background. For Batch 1 samples, we additionally separated the tissue in each organoid into two regions, which we called the inner and outer rings. These were clearly visually distinctive ring-shaped regions in the b=0 images (See Figure 1).

### Analysis strategy

The analyses were organized around the two questions posed in this study. First, to assess whether diffusion contrast captures reproducible spatial organization, we fitted DKI at each diffusion time, constructed normalized spatial radial profiles of MD, FA, and MK, and quantified inter-organoid agreement using Lin’s concordance correlation coefficient; DAPI staining provided a qualitative cytoarchitectural reference for the undirected batch. Second, to characterize the diffusion-time dependence of the signal and its relation to inter-compartment exchange, we quantified voxelwise MD changes with diffusion time, fitted NEXI, examined associations between DKI and NEXI parameters, and used analytical and Monte-Carlo two-compartment simulations to identify microstructural regimes capable of reproducing the observed apparent MD evolution. Finally, we performed a complementary analysis of Monte Carlo trajectories to compare displacement-derived diffusivity, the second-cumulant diffusivity of the PGSE signal, and finite-*b* ADC and DKI-derived MD estimates. The methodology and numerical validation of this analysis are detailed in Supplementary 7.1.

### Diffusion Signal Reconstructions

We applied Diffusion Kurtosis Imaging ^37;46^ (DKI) to both batches for each diffusion time. It was fitted to the dMRI signal using a weighted least squares algorithm implemented in the DIPY library ^47^. We extracted the following relevant parameter estimations for each organoid tissue voxel after fitting: Mean Diffusivity (MD), Fractional Anisotropy (FA), and Mean Kurtosis (MK).

The mean diffusivity estimated from DKI is interpreted here as a finite-*b* apparent quantity. For a normalized, direction-averaged diffusion signal, the cumulant expansion around *b* = 0 can be expressed as:

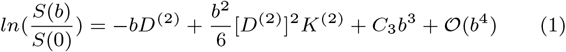

where *D*^(2)^ and *K*^(2)^ denote the second- and fourth-cumulant diffusion metrics, respectively. DKI models the signal up to the quadratic term, but higher-order contributions may influence the estimated MD and MK when fitting over a finite *b*-value range. Consequently, DKI-derived MD does not necessarily correspond to the second-cumulant diffusivity, *D*^(2)^ = −*∂*_*b*_ ln[*S*(*b*)*/S*(0)]|_*b*=0_. Throughout this study, MD therefore refers to the apparent mean diffusivity obtained from the specified DKI fitting window.

To select the DKI fitting range, we evaluated four candidate *b*- value windows (*b* ≤ 2000, 3500, 5000, and 7000 s/mm^2^) separately for each batch and diffusion time using two independent diagnostics. First, we quantified parameter underdetermination as the percentage of masked voxels yielding non-physical mean kurtosis values (MK *<* 0). Second, we quantified model misspecification as the largest systematic fit residual across shells, computed from powder- and voxel-averaged signals and normalized by the batch-specific noise level. Noise was estimated as the median coefficient of variation across the twelve repeated *b* = 0 volumes.

The two batches differed by approximately a factor of two in apparent diffusivity (mono-exponential ADC over *b* ≤ 1000 s/mm^2^: 0.59 and 0.30 *µ*m^2^/ms for Batch 1 and Batch 2, respectively), such that the same *b*-value corresponded to substantially different *bD* regimes. This difference is also evident in the powder-averaged signal decay shown in Supplementary Figure S1, Panel A. We therefore selected, for each batch, the largest *b*-window that maintained physically admissible kurtosis estimates while limiting systematic fitting errors relative to the batch-specific noise level. This procedure yielded *b* ≤ 3500 s/mm^2^ for Batch 1 and *b* ≤ 7000 s/mm^2^ for Batch 2. The corresponding diagnostics across all candidate windows are reported in Supplementary Figure S1.

To verify that the diffusion-time dependence is present in the signal itself, independently of the kurtosis fit and of the different *b*-windows used in the two batches, we additionally estimated a low-*b* ADC from the direction-averaged signal at *b* = 0 and *b* = 1000 s/mm^2^ alone, as 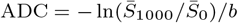, for each diffusion time and each batch. At this *b*-value, the signal decays by only 44% (Batch 1) and 26% (Batch 2), corresponding to *bD* ≈ 0.6 and 0.3. Although this estimate uses a lower *b*-value than the complete DKI fitting ranges, it remains a finite-*b* measurement and may still be influenced by non-Gaussian signal contributions.

The two-compartment Neurite Exchange Imaging model (NEXI ^3^) was fitted to both organoid batches. For the NEXI noise-floor correction, the noise standard deviation (*σ*) was estimated from the high-SNR *b* = 0 and *b* = 1000 s/mm^2^ data (*N* = 48 volumes/batch), and the signal was corrected for Rician bias before fitting ^48^. Data were powder-averaged and normalized by the mean value of the *b* = 0 s/mm^2^ volumes before NEXI fitting. NEXI fitting yielded the following parameter estimates per voxel: intra-neurite diffusivity (*D*_*i*_), extra-compartment diffusivity (*D*_*e*_), neurite fraction (*f*_*n*_), and exchange time (*t*_*ex*_).

We compared the parameter estimates of the two signal models at the voxel level to assess their consistency. For each organoid tissue voxel, the DKI-derived metrics (MD, FA, and MK) were related to the NEXI parameters (*D*_*i*_, *D*_*e*_, *f*_*n*_, and *t*_*ex*_) using Spearman’s rank correlation coefficient (*ρ*), which does not assume a linear relationship and is robust to outliers. Because MD and MK can vary with diffusion time, the correlations were computed both on the Δ-averaged DKI metrics and separately at each diffusion time (Δ = 15, 26, 38 ms). Correlations were computed across all tissue voxels, both separately for each batch and in the pooled data, so that a relationship was retained only when it was consistently signed across batches. We report correlation magnitudes rather than *p*-values given the large volume of voxels.

### Spatial Reproducibility Analysis

To assess the spatial organization of microstructural properties within individual organoids, a spatial radial profiling approach was applied to all DKI-derived parametric maps: MD, FA, and MK at each diffusion time (Δ = 15, 26, 38 ms).

We calculated the Euclidean distance from each voxel to the centroid and normalized it to obtain a radial distance in the range [0, 1] for each organoid in both batches, where 0 corresponds to the organoid center and 1 to its periphery. Normalized distances were divided into 10 equal-width bins spanning [0, 1], each representing 10% of the normalized radius. For each bin, the mean and standard deviation of the metric of interest were computed across all voxels falling within that shell.

The agreement between the radial profiles of all organoid pairs was measured using Lin’s concordance correlation coefficient (CCC) ^49^ to quantify the reproducibility of their radial organization descriptors within a batch. Unlike the Pearson or Spearman correlation, which are invariant to overall scale and offset and would therefore treat two profiles that share a trend but differ in absolute level as identical, CCC measures agreement with the line of identity, penalizing differences in both profile shape and absolute magnitude. It is a standard coefficient for reproducibility. Intra-batch reproducibility was defined as the mean CCC across the three pairwise organoid combinations within a batch (O1 vs O2, O1 vs O3, O2 vs O3), computed separately for each metric and diffusion time.

### Diffusion-time Dependence of MD

To characterize the diffusion time dependence of finite-*b* apparent MD at the voxel level, the DKI-MD parametric maps were independently calculated for each diffusion time (Δ = 15, 26, 38 ms) from both DKI fits (Batch 1: b ≤ 3500 s/mm^2^; Batch 2: b ≤ 7000 s/mm^2^). For each voxel within the tissue mask, a linear regression of MD against Δ was performed across the three diffusion times as a descriptive summary of the direction and magnitude of the MD change with Δ. The resulting slope, intercept, coefficient of determination (*R*^2^), and standard error of the slope were stored per voxel. We also correlated the NEXI-estimated parameters with MD slope values to capture, in a single correlation, how the MD increases (or decreases) with diffusion time could be affected by the more specific microstructural properties of the tissue estimated by NEXI.

### Simulations with the Two-Compartment Kärger Model

MD time-dependence was analyzed using a two-compartment Kärger-coupled two-compartment model with restricted spherical cells ^50^. The model comprises an intracellular compartment of restricted diffusion inside spheres of radius *R* with intrinsic diffusivity *D*_*i*_ (Gaussian-phase-distribution solution with reflecting boundaries), an extracellular compartment of Gaussian diffusion with diffusivity *D*_*e*_ and volume fraction 1 − *f* , and water exchange across the membrane described by the two-compartment Kärger model with permeability *κ*. The intracellular efflux rate follows the surface-to-volume ratio of the sphere, *k*_*i*_ = 3*κ/R*, and the influx rate from detailed balance, *k*_*e*_ = *k*_*i*_ *f/*(1 − *f*).

### Analytical parameter grid

Signals were computed on a grid of *R* ∈ {2, 3, 4, 5, 8, 10, 15} µm, *D*_*i*_, *D*_*e*_ ∈ {0.1, 0.2, 0.4, 0.6, 0.7, 1.0, 1.5} µm^2^/ms, *f* ∈ {0.25, 0.35, 0.5, 0.6, 0.65, 0.75} and *κ* ∈ {0, 10, 20, 25, 50, 80} µm/s, i.e. 12,348 substrates spanning impermeable to highly permeable cells, small to large cell bodies, and every ordering of intra- and extracellular diffusivity.

### Recreation of the full acquisition

The complete protocol was reproduced: *b* = 0, 1000, 2000, 3500, 5000, 7000 s/mm^2^, Δ = 15, 26, 38 ms, *δ* = 4.5 ms, and TE = 54 ms, using the measured gradient table (four *b* = 0 and 12/16/24/30/40 directions per shell, 126 volumes per diffusion time). Because the substrate is isotropic, every direction within a shell carries that shell’s signal, so the simulated data reproduced the acquisition’s number of measurements per shell, which determines each shell’s relative weight in the fit. The simulated volumes were then analysed exactly as the organoid data: a DKI fit (DIPY, WLS) per diffusion time, restricted to each batch’s *b*-window (*b* ≤ 3500 s/mm^2^ for Batch 1, *b* ≤ 7000 for Batch 2), from which MD was extracted. The diffusion-time dependence was summarised as ΔMD = MD(Δ=38)−MD(Δ=15). For comparison with a single-shell analysis, the shell-wise mono-exponential ADC was also computed. To relate the outcome to intracellular restriction, we defined the apparent intracellular diffusivity of the isolated, non-exchanging restricted-sphere signal as *D*_*i*,app_ = − ln *S*_sphere_*/b*, evaluated at *b* = 1000 s/mm^2^, Δ = 15 ms.

### Organoid regime and recreation of the measured curves

We identified the best-matching substrate for each batch by exhaustively searching all 12,348 simulated substrates and minimising the root-mean-square error (RMSE) between the simulated and measured tissue-averaged MD across the three diffusion times, evaluating each batch in its own DKI *b*-window. We did not constrain any parameter to its NEXI estimate because the two models assign different geometries to their intracellular compartments; their *D*_*i*_ and *f* are not one-to-one microscopic equivalents, and restricting the search by the NEXI values would not be justified. Fit quality was assessed against the standard error of the measured tissue-mean MD, and the parameter region compatible with the data was characterised from the best-ranked substrates.

### Monte-Carlo simulations

To test whether the positive diffusion-time dependence predicted by the analytical exchange model persists in an explicit cellular geometry, we selected the top ten substrates with the lowest RMSE between analytical and measured tissue-averaged MD at the three diffusion times for each batch (20 substrates in total), and we simulated 3D substrates of non-overlapping spheres using a Monte-Carlo random-walk simulator ^51^ (latest version available 2.2.0). The same parameters of the previous analytical simulations were retained for the Monte-Carlo ones. Each substrate consisted of spheres of radius *R* at intracellular volume fraction *f* in a voxel with side length 100 *µ*m, with intrinsic intracellular and extracellular diffusivities *D*_*i*_ and *D*_*e*_ and membrane permeability *κ*. One packing realization was used per parameter combination (Supplementary Table S1).

**Table S1.** Parameters and diffusion-time dependence of the twenty substrates selected from the analytical search for explicit Monte-Carlo simulation. Rank is defined by the analytical model’s RMSE against the measured tissue-averaged MD at three diffusion times, using each batch’s DKI fitting window. The parameters *R, f* , *D*_*i*_, *D*_*e*_, and *κ* were transferred to Monte Carlo without optimization. ΔMD is reported as percent change from 15 to 38 ms, separately for the Monte-Carlo (MC) and analytical models. Batch 1: *b* ≤ 3500 s/mm^2^; Batch 2: *b* ≤ 7000 s/mm^2^. All simulated substrates increased in MD over the sampled range.

| Batch | Rank | $R$ ($\mu\text{m}$) | $f$ | $D_i$ ($\mu\text{m}^2/\text{ms}$) | $D_e$ ($\mu\text{m}^2/\text{ms}$) | $\kappa$ ($\mu\text{m}/\text{s}$) | MC $\Delta\text{MD}$ (%) | Analytical $\Delta\text{MD}$ (%) |
| --- | --- | --- | --- | --- | --- | --- | --- | --- |
| 1 | 1 | 5 | 0.50 | 0.1 | 1.5 | 50 | +5.3 | +7.9 |
| 1 | 2 | 10 | 0.50 | 0.1 | 1.5 | 80 | +3.1 | +7.9 |
| 1 | 3 | 4 | 0.50 | 0.1 | 1.5 | 50 | +5.4 | +8.0 |
| 1 | 4 | 8 | 0.50 | 0.1 | 1.5 | 80 | +4.5 | +8.3 |
| 1 | 5 | 3 | 0.50 | 1.5 | 1.5 | 50 | +2.0 | +6.5 |
| 1 | 6 | 2 | 0.50 | 1.5 | 1.5 | 50 | +3.3 | +8.5 |
| 1 | 7 | 3 | 0.50 | 0.1 | 1.5 | 50 | +5.5 | +7.8 |
| 1 | 8 | 3 | 0.50 | 1 | 1.5 | 50 | +1.5 | +5.3 |
| 1 | 9 | 2 | 0.50 | 0.1 | 1.5 | 25 | +5.1 | +8.8 |
| 1 | 10 | 5 | 0.50 | 0.2 | 1.5 | 25 | +1.3 | +1.7 |
| 2 | 1 | 3 | 0.65 | 1 | 1.5 | 25 | +6.6 | +15.8 |
| 2 | 2 | 3 | 0.60 | 1.5 | 1.5 | 20 | +9.4 | +17.2 |
| 2 | 3 | 2 | 0.50 | 0.6 | 1 | 10 | +5.9 | +10.7 |
| 2 | 4 | 3 | 0.65 | 0.7 | 1.5 | 25 | +4.5 | +12.2 |
| 2 | 5 | 8 | 0.60 | 0.1 | 1.5 | 25 | +10.1 | +13.7 |
| 2 | 6 | 2 | 0.50 | 0.7 | 1 | 10 | +6.7 | +11.5 |
| 2 | 7 | 5 | 0.60 | 0.1 | 1.5 | 20 | +11.8 | +15.5 |
| 2 | 8 | 10 | 0.60 | 0.1 | 1 | 50 | +5.3 | +11.3 |
| 2 | 9 | 2 | 0.50 | 0.4 | 1 | 10 | +4.4 | +8.9 |
| 2 | 10 | 8 | 0.60 | 0.1 | 1 | 50 | +7.1 | +12.6 |

Monte-Carlo simulations used 10^6^ walkers propagated for 55 ms with 1000 time steps. The full experimental diffusion-encoding protocol was simulated at Δ = 15, 26, and 38 ms, with the measured gradient directions, *b*-values, and *δ* = 4.5 ms; *T*_2_ weighting was not included. The resulting signals were analyzed with the same DKI fitting procedure and batch-specific *b*-windows used for the organoid data (*b* ≤ 3500 s/mm^2^ for Batch 1 and *b* ≤ 7000 s/mm^2^ for Batch 2). For each substrate, the diffusion-time effect was summarized as ΔMD = MD(38 ms) − MD(15 ms), and the mono-exponential ADC between *b* = 0 and 1000 s/mm^2^ was calculated as a complementary finite-*b* estimate.

## Data Availability

The data supporting the findings of this study are currently being prepared for public release and will be made available in a public repository.

## Code availability

The code used in this study is currently being prepared for public release and will be made available in a public repository.

## Supplementary Material

Physical and finite-*b* apparent diffusivity in permeable cellular substrates

## Theoretical considerations

The finite-*b* mono-exponential ADC differs from the second-cumulant diffusivity of the diffusion signal. A mono-exponential ADC measured at a finite *b*-value is

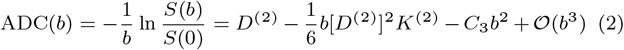

where *D*^(2)^ is the second-cumulant diffusivity, *K*^(2)^ is the corresponding dimensionless kurtosis in the *b* → 0 limit, and *C*_3_ denotes the coefficient of the cubic term in the log-signal expansion. Then a finite-*b* ADC is an apparent diffusion coefficient and does not generally equal *D*^(2)^.

Independently of the preceding low-b cumulant expansion, we now consider a non-exchanging mixture of compartmental signals with fixed signal fractions. The following result holds at arbitrary *b* and does not require Gaussian diffusion within individual compartments. Let *d*_*j*_ (*b*, Δ) = − ln[*S*_*j*_ (*b*, Δ)*/S*_*j*_ (0)]*/b* denote the finite-*b* ADC of compartment *j*, with fixed positive signal fractions *w*_*j*_ satisfying ∑_*j*_ *w*_*j*_ = 1. The normalized mixture signal is then 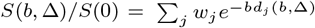. Differentiating the mixture ADC at fixed *b* gives

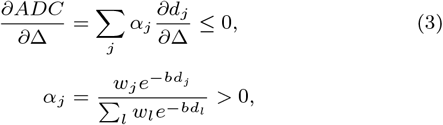

provided that *∂d*_*j*_*/∂*Δ ≤ 0 for every compartment. Therefore, for non-exchanging compartments with fixed signal fractions, the mixture ADC cannot increase with Δ if each compartment’s ADC is non-increasing.

**Figure S1.**
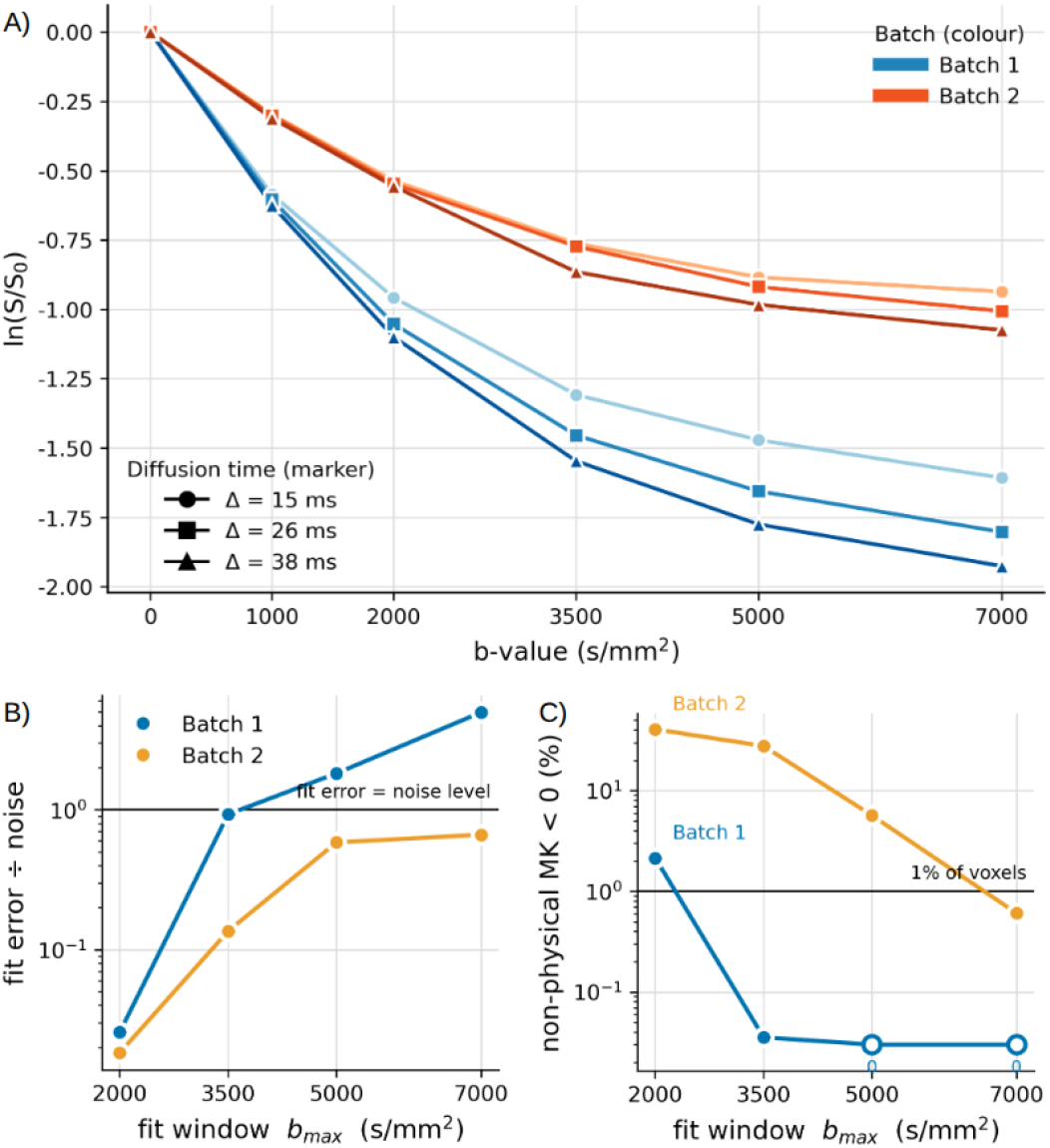
Batch-specific DKI fit-window selection. **Panel A)** Powder-averaged signal decay for Batch 1 (blue) and Batch 2 (orange), with the three diffusion times (Δ = 15, 26, 38 ms) overlaid per batch (markers). Batch 2 decays less steeply than Batch 1 across all shells, consistent with its approximately two times lower apparent diffusivity (mono-exponential ADC over b *≤* 1000 s/mm^2^: 0.59 vs. 0.30 µm^2^/ms). **Panel B)** Worst systematic fit error across shells, normalised by each batch’s own noise level (defined as the median coefficient of variation of the twelve repeated b = 0 volumes). **Panel C)** Parameter determinacy: percentage of masked voxels returning non-physical MK *<* 0; open markers indicate 0%. The horizontal reference lines mark the 1% of non-physical voxels (C); and the fit error = 1× noise (B).

In contrast, when exchange is introduced through the stationary two-Gaussian-compartment Kärger model, 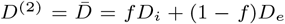 is independent of exchange rate and diffusion time, whereas

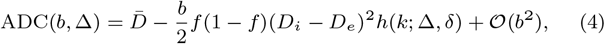

where *k* = *k*_*i*_ + *k*_*e*_ is the total exchange rate and *f* is the intracellular fraction. The dimensionless exchange-weighting factor is defined as 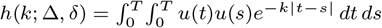, where *u*(*t*) = |**q**(*t*)|^2^*/b* is the normalized diffusion-encoding weight, with *b* = 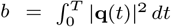. Here, 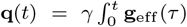 *dτ* is the time-dependent accumulated dephasing wavevector, and **g**_eff_ (*t*) is the effective diffusion-encoding gradient waveform, accounting for the refocusing pulse.

In the stationary two-compartment Kärger model, exchange allows water molecules to sample different compartmental diffusivities during the PGSE encoding period (motional averaging). As diffusion time increases, the factor *h* and the corresponding second-cumulant kurtosis decrease, allowing the finite-*b* ADC to increase without any change in *D*^(2)^.

**Figure S2.**
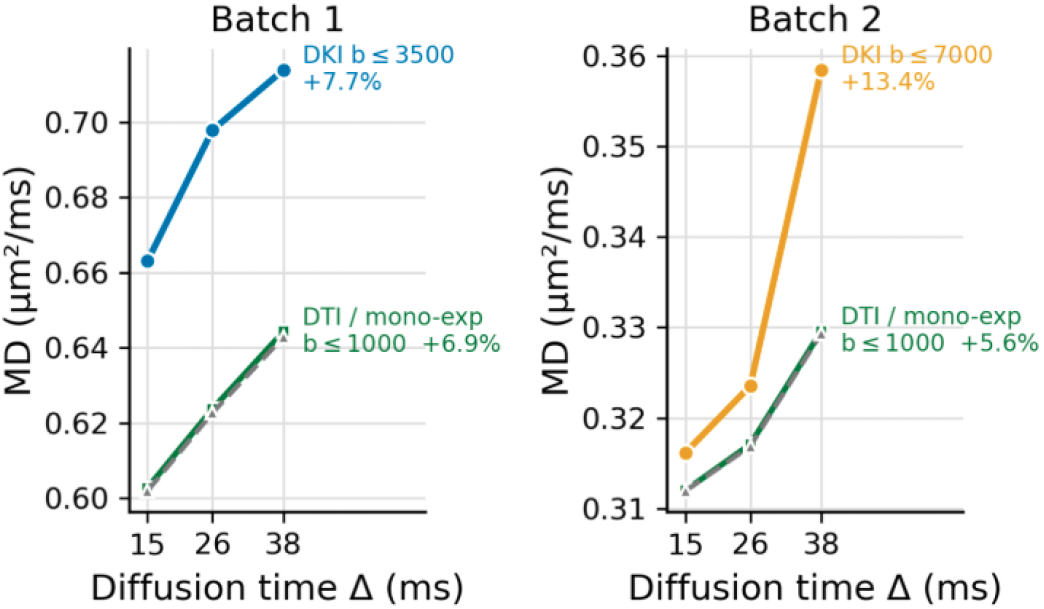
Batch-averaged diffusivity against diffusion time. Grey dashed: low-*b* ADC from the direction-averaged signal at *b* = 0 and *b* = 1000 s/mm^2^ alone, 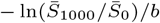; green: tensor fit over the same two shells; blue/orange: kurtosis fit over each batch’s full *b*-window (*b ≤* 3500 and *b ≤* 7000 s/mm^2^). Labels give the change between Δ = 15 and 38 ms. The two low-*b* estimators agree within 0.3%, and both show increasing apparent diffusivity in the two batches, although these finite-*b* estimates may still contain appreciable non-Gaussian contributions. Means over all tissue voxels (*n* = 5,643 and 9,038).

**Figure S3.**
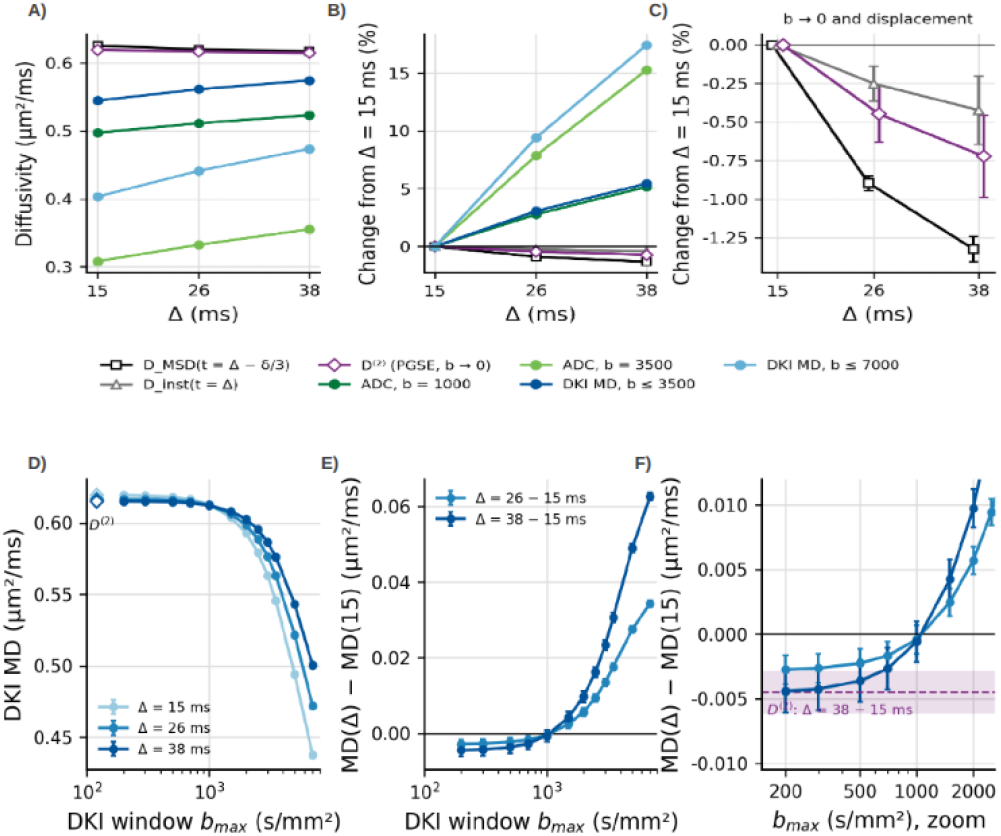
Physical and finite-*b* apparent diffusivity in a permeable Monte Carlo substrate. Results for the best-matching Batch 1 substrate (*R* = 5 µm, *f* = 0.5, *D*_*i*_ = 0.1, *D*_*e*_ = 1.5 µm^2^/ms, *κ* = 50 µm/s), evaluated after equilibration. Error bars and shaded intervals represent 95% confidence intervals estimated using a paired jackknife. **(A)** Diffusion-time dependence of displacement-derived diffusivity (*D*_MSD_; black), instantaneous diffusivity (*D*_inst_; gray), PGSE second-cumulant diffusivity (*D*^(2)^, *b →* 0; purple), mono-exponential ADC at *b* = 1000 (dark green) and 3500 s/mm^2^ (light green), and DKI-derived MD fitted over *b* ≤ 3500 (dark blue) and 7000 s/mm^2^ (light blue). **(B)** Relative changes in the same estimators with respect to Δ = 15 ms. **(C)** Enlarged view highlighting the decreasing physical and second-cumulant diffusivities. **(D)** DKI-derived MD as a function of the maximum *b*-value included in the fit (*b*_max_), for each diffusion time (15ms - light blue; 26ms - blue; 38ms - dark blue). **(E)** Differences in fitted DKI-MD between Δ = 26 (blue) or 38 ms (dark blue) with Δ = 15 ms across fitting windows. **(F)** Enlarged low-*b* region showing the reversal from decreasing to increasing fitted MD, with the second-cumulant difference indicated by the purple dashed line and shaded interval.

This reference model assumes Gaussian compartments and does not describe restricted intracellular motion. Moreover, a decrease in kurtosis alone does not explain an increase in DKI-fitted MD since DKI already models the quadratic term in *b*. So for noise-free signals, differences between finite-window DKI MD and *D*^(2)^ involve neglected terms of order *b*^3^ and above in the log-signal.

## Trajectory-based validation

We revisited the best-matching Batch 1 substrate using explicit Monte-Carlo trajectories: a periodic 100-*µ*m cube with non-overlapping spheres (*R* = 5 *µ*m, *f* = 0.5, *D*_*i*_ = 0.1 and *D*_*e*_ = 1.5 *µ*m^2^/ms, *κ* = 50 *µ*m/s). One million walkers from 20 independently seeded runs were propagated for 220 ms (*dt* = 0.055 ms); PGSE encoding (*δ* = 4.5 ms, Δ = 15, 26, 38 ms) began after 165 ms of equilibration. We measured

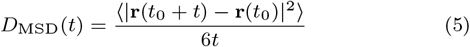

Displacement-based diffusivities were calculated from unwrapped trajectories, averaging MSD over 11 late time origins (110–165 ms), where displacement statistics were approximately stationary. For comparison with PGSE, *D*_MSD_ was evaluated at *t* = Δ − *δ/*3, and *D*_inst_ = (1*/*6)*d*MSD*/dt* at *t* = Δ.

The waveform-specific second-cumulant diffusion coefficient was independently obtained from the accumulated phase covariance, our findings more effectively.

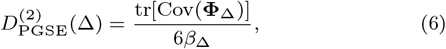

where **Φ**_Δ_ is the three-dimensional phase per unit gradient amplitude and *β*_Δ_ is the corresponding waveform *b*-value per unit squared amplitude. From these trajectories, we estimated finite-*b* ADC at *b* = 1000 and 3500 s/mm^2^, and DKI-derived MD using the *b* ≤ 3500 and 7000 s/mm^2^ fitting windows. Also, other densely sampled synthetic *b*-windows were used to investigate the convergence of fitted MD towards the small-*b* limit. We repeated the analysis with impermeable membranes (*κ* = 0) in the same geometry. Uncertainties on paired diffusion-time differences were estimated by a 40-group jackknife over walkers.

## Results and numerical controls

Between Δ = 15 and 38 ms, the permeable substrate showed decreasing *D*_MSD_(*t* = Δ − *δ/*3) (−0.0083 *µ*m^2^/ms; 95% CI [−0.0088, −0.0078]), *D*_inst_(*t* = Δ) (−0.0026; [−0.0040, −0.0012]), and 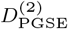 (−0.0045; [−0.0061, −0.0028]), while ADC at *b* =1000 s/mm^2^ increased (+0.0257; [+0.0246, +0.0267]), as did DKI-derived MD at *b* ≤ 3500 s/mm^2^ (+0.0297; [+0.0284, +0.0309]; Supplementary Fig. S3, Panels A-B-C). The small-*b* kurtosis decreased from 2.05 to 1.52. The ADC ordering changed near *b* = 129 ± 45 s/mm^2^, while the fitted DKI MD difference approached zero near *b*_max_ = 1000 s/mm^2^ and became negative at smaller windows (Supplementary Fig. S3, Panels D-E-F). In the impermeable control, all five diffusivity estimates decreased with diffusion time, and the small-*b* kurtosis showed no resolved change. The sign reversal persisted across start times, time steps, and two alternative sphere packings. Thus, exchange can increase finite-*b* apparent diffusivity while physical and second-cumulant diffusivities decrease in this substrate. This establishes a sufficient mechanism, but not exchange specificity or the physical diffusivity of the organoids.

## Acknowledgments

This work was supported by the Swiss National Science Foundation 320030-2277717 to EFG.

E.J.C.R. acknowledges financial support from the “Ramón y Cajal” Excellence Fellowship (Grant RYC2023-042763-I), funded by the Ministry of Science, Innovation and Universities (MICIU) and the Spanish State Research Agency (AEI, 10.13039/501100011033), and co-financed by the European Social Fund Plus (ESF+)

We thank Prof. Sune N. Jespersen for his insightful comments and constructive discussions, which prompted further investigation into the distinction between physical and finite-*b* apparent diffusivity and helped refine the interpretation of our findings.

We thank Prof. Oscar Esteban for his valuable feedback and constructive suggestions, which helped strengthen the manuscript’s narrative, improve the clarity of its presentation, and communicate

